# IL1β/IL1R1/IRAK4 Drives Inflammatory Ovarian Cancer Seeding at the inflamed sites and Is Reversed by an IRAK4 inhibitor UR241-2

**DOI:** 10.64898/2026.04.30.722105

**Authors:** John P. Miller, Kyu Kwang Kim, Cameron WA Snyder, Negar Khazan, Niloy A. Singh, Megan E. Boyer, Elizabeth Lamere, Myla Strawderman, Sonali Sharma, Ronald Lakony, Michelle Whittum, Mark Anderson, Rick Keenan, Elizabeth Pritchett, Cameron Baker, John Ashton, Manoj K. Khera, Michael R. Elliott, Christina M. Annunziata, Jeevisha Bajaj, Laura M. Calvi, Michael W. Becker, Rachael Rowswell-Turner, Richard G. Moore, Rakesh K. Singh

## Abstract

Inflammation-driven tumor implantation, such as port-site metastasis (PSM) following laparoscopic gynecologic surgery and peritoneal seeding during post-surgical recurrence, represents an aggressive clinical problem that remains poorly understood and lacks targeted therapies. To address this, we developed a non-surgical Mesothelium-Inflammation/Injury-Metastasis (MIM) model and investigated the role of the IL-1β/IL1R1/MYD88/IRAK1/4 axis and NLRP3 in epithelial ovarian cancer (EOC) seeding at inflamed or injured sites. This model created by a needle injury recapitulates inflammation-driven peritoneal seeding and mimics PSM and inflammation associated dissemination in peritoneum during recurrence. Seeding was dependent on Il1r1 but not Nlrp3, despite its role in regulating IL-1β production, as Il1ra⁻/⁻ and Nlrp3⁻/⁻ mice phenocopied wild-type C57BL/6 mice. Given the limited antitumor efficacy of IL-1β–targeting agents such as Anakinra and Canakinumab, we focused on IRAK4 as a therapeutic target. IRAK4 knockdown significantly prolonged survival, reduced tumor cell adhesion, downregulated E-cadherin and Wnt4, and induced S-phase/mitotic arrest. This led to the development of UR241-2, a small-molecule IRAK4 inhibitor, which was validated through molecular simulations, hotspot analysis, nanoBRET, global kinome profiling, and NF-κβ reporter assays. UR241-2 inhibited NF-κβ nuclear translocation and blocked IL-1β–induced IRAK4 phosphorylation. UR241-2 exhibited favorable drug-like properties, including absence of CYP or hERG inhibition, and acceptable CaCo-2 permeability, plasma protein binding, microsomal stability, and pharmacokinetics. In vivo, UR241-2 reduced SKOV3 xenograft growth, suppressed mesothelial seeding, and increased MHC-II⁺ macrophages and activated neutrophils in syngeneic high-grade epithelial ovarian HGS3 tumors. RNA-seq revealed enrichment of neutrophil activation signatures and suppression of extracellular matrix (ECM) gene programs. Together, these findings establish a role for the IL-1β/IL1R1/IRAK4 axis in inflammation-driven PSM and peritoneal seeding and ECM regulation in EOC, and demonstrate that IRAK4 inhibition activates antitumor immune responses, providing a therapeutic strategy to block metastatic seeding and improve tumor control.

## Introduction

Chronic inflammation (CI) is a pathological feature of epithelial ovarian cancer (EOC)^1–2^ and most other malignancies^3^. A clear clinical manifestation of inflammation-driven tumor seeding is port-site metastasis (PSM) following laparoscopic gynecologic surgery, which often reflects aggressive or advanced disease^4–5^. Similar metastases are observed in other cancers, including catheter-site seeding in hepatocellular carcinoma^6^ and tumor implantation at surgical anastomotic sites in gastric cancer. Together, these observations highlight that tumor seeding at sites of tissue injury and inflammation is a clinically significant and widespread problem. However, despite its importance, this clinical event remains poorly understood and lacks targeted therapeutic strategies. As a result, PSM and other inflammation-related metastatic seeding events in EOC and other risky malignancies continue to be managed with radiation, surgery, or cytotoxic chemotherapy rather than targeted therapies. Chronic inflammation recruits and sustains activation of the IL1β/TLR/IRAK4/NF-κβ axis^7^. In EOC, each of the key components of this pathway: including IL1β^8–10^, its receptor IL1R^11–12^, the myddosome complex partners TLR4/Myd88^13–20^ and NF-κβ^21–30^ are aberrantly overexpressed, predicting poor survival and chemoresistance^31^. IL1β, in particular, has been implicated in multiple steps of peritoneal dissemination^32^, including loss of mesothelial integrity^33–34^, formation of solid peritoneal tumors from ascitic cells, and enrichment of pro-tumorigenic cytokines in the peritoneal microenvironment^32, 35–39^.

Despite the importance of this inflammatory axis in EOC and other cancers, agents that directly target IL-1 or its receptor IL1R, such as Anakinra and Canakinumab⁴⁰ have not demonstrated clear therapeutic benefits in the malignancies tested. Similarly, targeting MYD88, a key downstream component of IL1R1 signaling, has not demonstrated clinical benefits in EOC. Therefore, we examined whether IRAK1 and IRAK4, the central kinases in IL-1β signaling and key downstream effectors of IL1R1/MYD88, can offer the desired therapeutic responses. Notably, selectively targeting IRAK1 was shown to attenuate hyaluronic acid-induced stemness and non-canonical STAT3 activation in EOC^41^. Further, TCS2210, a selective IRAK1 inhibitor, also suppressed EOC growth in vivo^41^, supporting the therapeutic promise of targeting IRAK1/IRAK4 in EOC. However, the roles of IRAK4^42–43^, the master regulator of IRAK1 and the key mediator of the myddsome complex signaling remain largely unexplored in EOC, and no targeted therapies against IRAK4 are currently being tested for EOC treatment.

Here, we investigated the expression of IRAK4 in EOC tissues and examined the role of IRAK4 in EOC cell adhesion to the peritoneal mesothelium, as a model site for injury induced inflammation and its contribution to immune dysfunction, alongside IL1β/IL1R1 and NLRP3 signaling. We developed a mechanical injury–induced Mesothelium Inflammation (MiM) model that recapitulates inflammation-driven peritoneal seeding in-vivo and post-surgical PSMs in EOC and other malignancies. This model enabled direct evaluation of IL1R1, its antagonist IL1RN, and NLRP3 in regulating tumor seeding and burden, and provided a platform to test current or future IRAK4-targeted therapies for anti-adhesion activity in EOC or other malignancies. Using structure–activity relationship–guided medicinal chemistry, we developed UR241-2, a novel small-molecule IRAK4 inhibitor, and evaluated its activity in vitro and in vivo. Comprehensive profiling, including kinome selectivity, comparison with clinical IRAK4 inhibitors (PF-06650833, CA4948), pacritinib, CTX-204885, bosutinib, and CEP-701, as well as pharmacokinetic and ADME analyses, demonstrated favorable drug-like properties. Notably, UR241-2 showed improved selectivity over CA4948, which exhibits off-target activity against FLT3 and CLK kinases, a counterproductive effect since high expression of these kinases is associated with improved outcomes in EOC. Efficacy studies in xenograft and syngeneic MiM models, combined with flow cytometry and RNA-seq, revealed that targeting IRAK4 reduces tumor seeding and modulates the tumor-immune microenvironment, specifically via activation of MHCII in macrophages and neutrophils. Taken together, these findings establish a role for the IL-1β/IL1R1/IRAK4 axis in inflammation-driven peritoneal seeding and ECM regulation in EOC, and demonstrate that IRAK4 inhibition activates antitumor immune responses, providing a novel therapeutic strategy to block metastatic seeding and improve tumor control.

## Results

### Il1β/Il1r1 loss reduces seeding of murine high-grade serous EOC at injury sites: development of the MiM model

Analysis of EOC patient microarray data showed that high IL-1β and IL-1R1 expression is associated with poor prognosis (Fig-1A-B). To investigate the role of IL-1β/IL-1R1 signaling in metastasis, we developed a murine Mesothelium-Inflammation/Injury-Metastasis (MiM) model (see Methods). HGS-3 murine high-grade EOC cells (3.7 × 10⁶ cells/mouse) were injected intraperitoneally into C57BL/6 mice using a 21-gauge needle. Tumor nodules formed at the site of needle injury within 30–45 days. To assess the role of IL1R1, HGS-3 cells were implanted into wild-type (WT) and *Il1r1* knockout (Il1r1KO; B6.129S7-Il1r1tm1Imx/J) mice (Fig-1C). After 42 days, tumors were collected from injury sites and omentum. WT mice developed cutaneous tumors in 80% of cases (Fig-1D), whereas Il1r1^KO^ mice showed reduced tumor incidence and smaller tumors at injury sites (Fig-1E). In contrast, omental tumor burden was similar between groups (Fig-1F). Tumor weights at inflamed cutaneous sites were significantly reduced in Il1r1^KO^ mice (Fig-1G). To further assess IL-1 signaling, we examined Il1rn knockout (Il1rn^KO^) mice. WT and Il1rn^KO^ mice^44^ developed comparable omental (Fig-1I) and cutaneous tumors (Fig-1L), with no significant differences in tumor size (Fig-1J, M). We next tested whether Nlrp3, a key inflammasome component critical in the activation of IL-1, contributes to tumor seeding. HGS-3 cells implanted in Nlrp3^KO^ (B6.129S6-Nlrp3tm1Bhk/J) mice showed tumor burden comparable to WT controls at both cutaneous and omental sites (Supplementary Fig-1). Immune profiling of peritoneal lavage and tumor tissue by flow cytometry showed no significant differences between WT and Il1r1^-/-^ mice in F4/80⁺MHCII^hi/lo^ macrophages or CD11b⁺Ly6⁺ neutrophils (Fig-1M). These findings are consistent with the limited clinical efficacy of IL-1–targeting agents such as Anakinra and Canakinumab in cancer (Fig-1O). Similarly, no therapies targeting MYD88 or TLR pathways are approved for EOC or other malignancies (Fig-1O). We, therefore, focused on IRAK4, a central kinase downstream of IL-1R/TLR signaling (Fig-1P and Fig-1Q), which is currently under investigation in multiple clinical trials including NCT07064122 (treatment of hematologic malignancies), NCT05669352 (treatment of brain metastasized melanoma), CT06696768 (treatment of colorectal cancer) and NCT04933799 (Covid treatment), NCT07137637 (treatment of relapsed and refractory acute myeloid leukemia).

**Figure-1.**
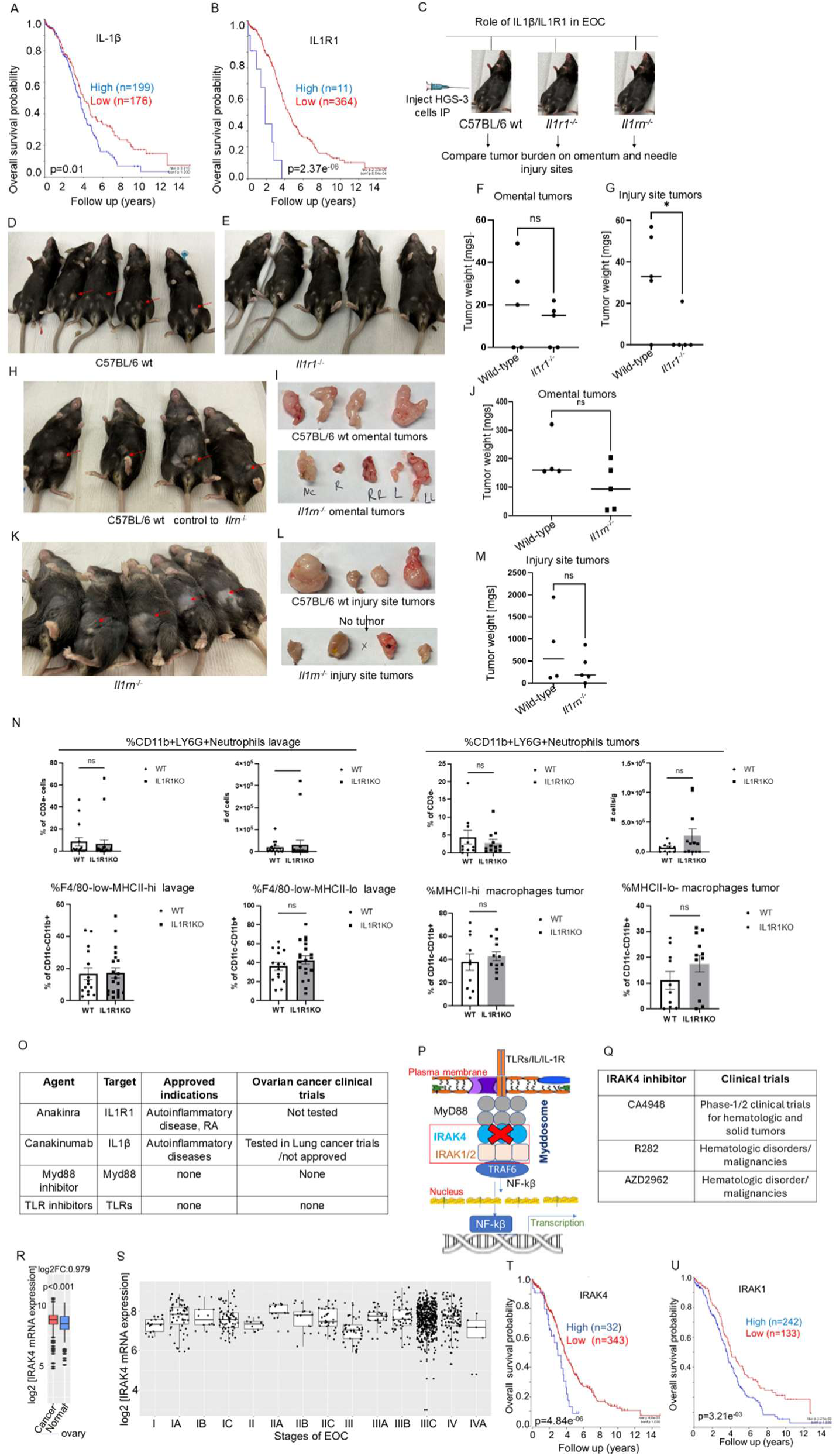
: IL1R1 is central to adhesion of EOC cells on the inflamed peritoneal walls in mice. (**A-B**): Analysis of microarray data (TCGA-381-tpm-gencode36) of ovarian cancer patients using R2 Genomics Analysis and Visualization Platform tools showed that IL1β and IL1R1 mRNA overexpression predicts poor survival. (**C**): Schema of testing the role of Il1β, Il1r1 and Il1rn (Il1r1 antagonist) in ovarian cancer burden both at the needle injury and omental sites. (**D-E**): HGS-3 murine high-grade serous EOC cells (4.5 million/per mice) were implanted intraperitoneally using 21 sterile gauge needle in C57BL/6 WT and C57BL/6 *Il1r1*^KO^ mice. Mice were observed for 45-50 days and euthanized. Tumors formed on needle injury site both protruding at the skin, and, in the peritoneum, and on the omentum, shown by red arrows, were isolated, weighed and frozen in liquid nitrogen. Lavages via washing with sterile PBS(5mL) were also collected. The studies were repeated thrice. A representative experiment is shown. (**F**): Weights of the omental did not differ between C57BL/6 wt and *Il1r1*^KO^ mice. (**G**): Weights of the peritoneal/skin tumors differed significantly between C57BL/6 wt and *Il1r1*^KO^ mice groups. * indicates <0.05. (**H-K**): Tumors formed at the needle injury site in C57BL/6 wt and *Il1rn*^KO^ mice are pointed with the red arrows. (**I**-upper vs **I**-lower): Images of the omental and peritoneal /skin tumors harvested from the euthanized *Il1r1*^KO^ mice are shown. (**J**): Omental tumor weights in C57BL/6 wt did not differ between *Il1rn^k^*° mice groups. (**L**-upper vs **L**-lower): Images of the omental and peritoneal /skin tumors harvested from the euthanized *Il1rn*^KO^ mice are shown. (**M**): Weights of the tumors formed on the needle injury site in C57BL/6 WT mice did not differ for *Il1rn*^KO^ mice groups. This experiment was repeated twice. Tumor sizes were analyzed via non-parametric T-test using Graph-Prism version -7 or higher. * indicates <0.05. (**N**): Il1r1 KO does not significantly impact myeloid percentages in the peritoneal lavage and tumor. A) Peritoneal lavage and tumor percentages and cell numbers of CD11b+Ly6G+ neutrophils from flow cytometry data of wild type and IL1R1KO mice injected with HGS3 tumors. B) Peritoneal lavage percentages and cell numbers of F4/80loMHCIIhi and F4/80hiMHCIILow macrophages from the peritoneal lavage and percentage of MHCII^hi^ and MHCII^Low^ macrophages from tumors from flow cytometry data of wild type and IL1R1 KO mice injected with HGS3 tumors. N = 15-20 mice per group from 3 independent experiments. Statistical significance was determined using Mann-Whitney test, *p < 0.033, **p < 0.002, ***p < 0.001. (**O**): Table shows that none of the IL1β, IL1R1 and myd88/TLR targeted agents has been approved for treatment of a malignancy yet. (**P**): Schema of IL1β/IRAK4 signaling pathway. Scheme shows that inhibiting IRAK4 can centrally block IL1/TLR driven signaling in ovarian malignancies. (**Q**): Table summarizes the status of 3 IRAK4 inhibitors undergoing clinical trials. (**R**): Using GENT2 database we observed that compared to normal ovaries, malignant ovaries overexpress IRAK4 mRNA. (**S**): GENT2 database also showed that higher stages of EOC disease significantly increased IRAK4 mRNA expression. Analysis of the EOC patient’s microarray data using GENT2 tools showed that IRAK4 mRNA expression was altered in various stages of disease. Two-sample T-test showed statistical differentiations: IA vs I (p=0.004); IA vs II (p=0.007); IA vs III (p<0.001); IC vs IIA (p=0.002); IC vs III (p<0.001); IIA vs I (p<0.001); IIA vs II (p<0.001); IIA vs III (p<0.001); IIA vs IIC (p=0.006); IIA vs IIIC (p=0.001); IIC vs III (p<0.001); III vs IIB (p=0.004); IIIA vs I (p=0.001); IIIA vs II (p=0.004); IIIA vs III (p<0.001); IIB vs I (p=0.008); IIIB vs III (p=<0.001); IV vs I (p=0.004); IV vs IIA (p=0.005); IV vs III (p=<0.001). (http://gent2.appex.kr/gent2/, date accessed 10/2/2023). (**T-U**): Analysis of microarray data (TCGA-381-tpm-gencode36) of ovarian cancer patients using R2 Genomics Analysis and Visualization Platform tools showed that IRAK4 and IRAK1 mRNA overexpression predicts poor survival in EOC patients.

### IRAK4 expression increases with disease stage and is associated with poor prognosis in EOC

Analysis of EOC patient microarray data using GENT2 tools^45^ showed that IRAK4 mRNA is significantly overexpressed in malignant ovarian tissue compared to normal ovary (p < 0.001, log2Fold change = 0.979) (Fig-1R). Stage-stratified analysis revealed that IRAK4 expression varies significantly across disease stages (Fig-1S; http://gent2.appex.kr/gent2/, accessed 10/2/2023). Pairwise comparisons showed consistent increases from early to advanced stages, including IA vs I (p = 0.004), IA vs II (p = 0.007), and IA vs III (p < 0.001). Significant differences were also observed between intermediate and advanced stages, including IC vs IIA (p = 0.002) and IC vs III (p < 0.001). Comparisons involving stage IIA showed strong differences across multiple groups (IIA vs I, II, III: all p < 0.001; IIA vs IIC: p = 0.006; IIA vs IIIC: p = 0.001). Additional differences were observed between IIC vs III (p < 0.001), III vs IIB (p = 0.004), IIIA vs I (p = 0.001), IIIA vs II (p = 0.004), and IIIA vs III (p < 0.001). Late-stage comparisons also remained significant, including IV vs I (p = 0.004), IV vs IIA (p = 0.005), and IV vs III (p < 0.001).

Independent analysis using the R2 Genomics platform confirmed that high IRAK pathway expression is associated with poor prognosis, including IRAK1 (p = 3.21 × 10⁻^3^) and IRAK4 (p = 4.8 × 10⁻^6^), while IRAK2 showed a weaker association (p = 0.062) (Fig-1T-U).

### IRAK4 knockdown enhances survival and suppresses adhesion and cell-cycle progression

IRAK4 shRNA knockdown in high grade serous EOC HGS-3 cells (Fig-2A) significantly improved survival in tumor-bearing mice compared to controls (Fig-2B) when implanted intraperitoneally in C57BL/6 mice. Proteomic analysis revealed increased WNT4 expression in control cells relative to IRAK4 knockdown cells (Fig-2C), consistent with its role as a poor prognostic marker.

Functionally, IRAK4 knockdown induced S-phase cell cycle arrest (Fig-2D), reduced mitotic progression (Fig-2E), and significantly decreased adhesion to collagen (Fig-2F). E-cadherin expression was also reduced (Fig-2G), and its expression positively correlated with IRAK4 levels in EOC tumors (Fig-2H).

**Figure-2:**
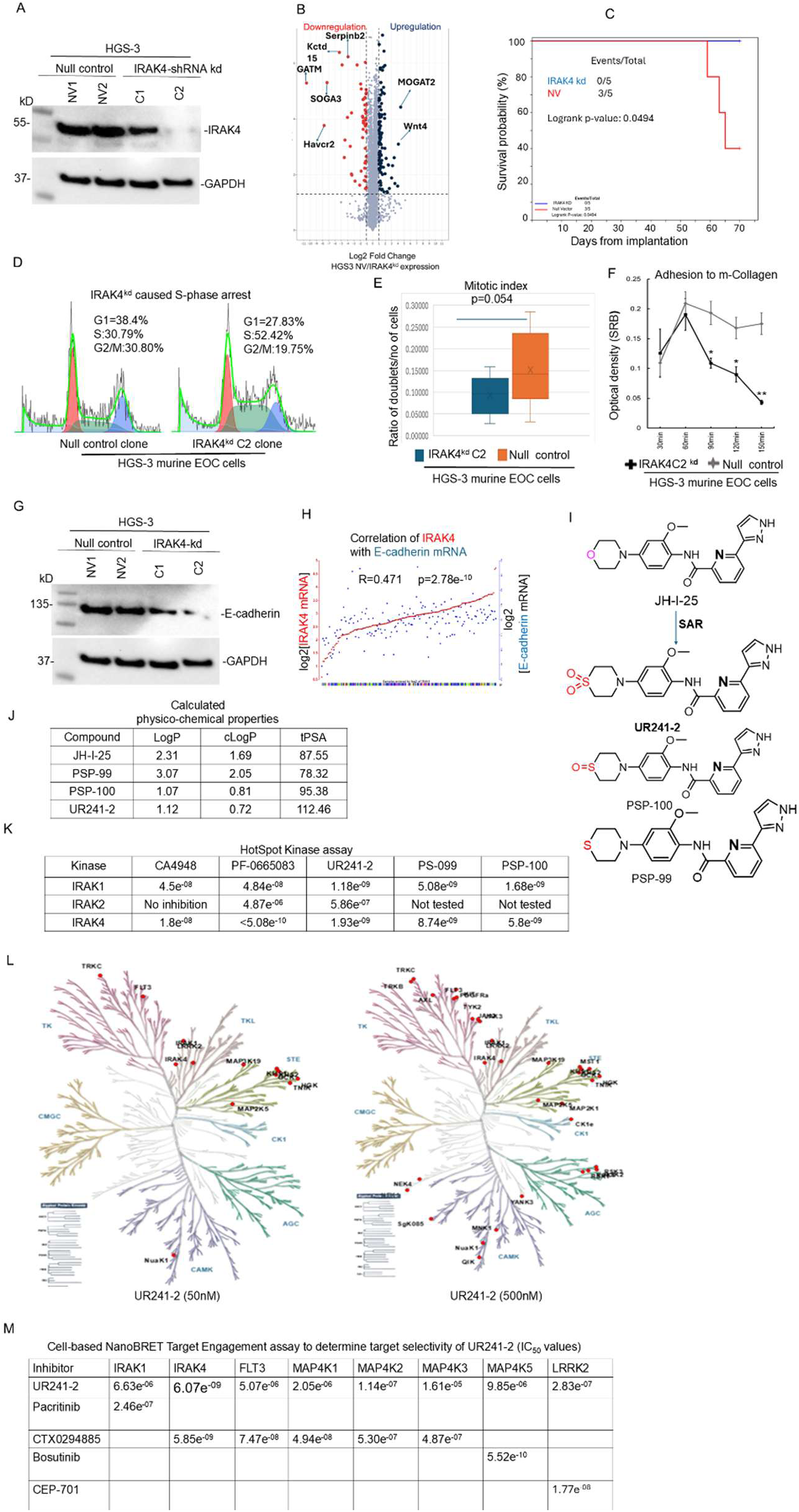
IRAK4 participates in survival, cell-cycle progression and cell adhesion. (**A**): IRAK4shRNA or null shRNA knockdown followed by clonal selection of HGS-3 murine EOC cells generated a C2-knockdown clone that showed nearly complete deletion of IRAK4. Null vector clones NV-1 or NV-2 were also generated following same procedures. (**B**): Proteomics analysis of the HGS-3 null vector NV vs partial IRAK4^kd^ showed very limited off-target effects. Notably, downregulation of WNT-4, a poor prognostic factor of EOC was observed. **(C)** : C2-IRAK4^kd^ HGS-2 clone cells (7 million/mice, IP) when implanted in C57BL/6 mice showed significant increase in survival than the NV1 null vector implanted mice (log rank p=0.0494). **(D)** : IRAK4^kd^ reduced S-phase arrest. S-phase population upon partial IRAK4^kd^ rose from 30.79 to 52.42% in HGS-3 EOC cells. **(E)** : IRAK4^kd^ reduced mitotic index in HGS-3 murine EOC cells (p=0.054). **(F)** : IRAK4^kd^ reduced adhesion to murine-Collagen. p=0.0071, p=0.0105, p=0.0009, error bars: one stdev in either direction. **(G)** : IRAK4^kd^ HGS clones resulted in reduced expression of E-cadherin. **(H)** : E-cadherin and IRAK4 mRNA show strong correlation of expression. R=0.471, p=2.7e^-10^. mRNA expression data available at R2-genomics was analyzed using the system inbuilt tools **(I)** : Structure-activity relationship (SAR) guided optimization of JH-I-25 scaffold, a literature described IRAK4 inhibitor leading to 3 potent novel analogs, is shown. The chemical structures of UR241-2 (sulfone), PSP-099 (sulfide) and PSP-100 (sulfoxide) are shown. **(J)** : Calculated drug-likeness/physicochemical properties (LogP, cLogP and topological polar surface area-tPSA are shown. These properties were calculated using Chemdraw software. **(K)** : Comparison of IRAK1, IRAK2, and IRAK4-kinase inhibitory IC_50s_ of UR241-2 versus CA4948, PF-06650833, PSP-099 and PSP-100 are shown. Compounds were screened using HotSpot Kinase assay available in Reaction Biology Laboratories using 1μM ATP concentration under a 10-dose singlet screening program. **(L)** : Dendrograms of global kinome activity of UR241-2 at 50- and 500nM doses. Red indicates kinase affected. At 50nM dose, 13 of 682 kinases were inhibited, whereas increasing dose to 500nM, 34 kinases, shown as red dots, from among 682 total kinases, were inhibited. **(M)** : A HEK293 cell-based nanoBret target Engagement screening assay was conducted to determine the selectivity of UR241-2 among 10 most affected kinases revealed by HotSpot kinase assay. The nanoBret assay showed that UR241-2 inhibits IRAK4 kinase activity selectively and other kinases including IRAK1 are affected at 10-100 folds higher doses, qualifying UR241-2 as one of the selective and specific IRAK4 kinase inhibitors.

### UR241-2 is a potent and selective IRAK4 inhibitor with minimal off-target activity

Given the role of IL-1/TLR/MYD88/IRAK4 signaling in EOC, we sought to develop a selective IRAK4 inhibitor. Leading compound, CA4948, currently in multiple clinical trials for AML and solid tumors, exhibited off-target activity involving three IRAK4/FLT3/CLK kinases. Notably, increased expression of CLK and FLT3 correlates with improved prognosis in EOC (Supplementary Fig-2), highlighting the need for improved selectivity.

We optimized the JH-I-25 scaffold, a reported IRAK4 inhibitor (∼20 nM potency), using bio-isosteric substitution of the morpholine moiety with thiomorpholine, thiomorpholine-S-oxide, and thiomorpholine-S-dioxide (Fig-2I). These modifications substantially improved IRAK4 IC₅₀ values and enhanced predicted physicochemical properties, including cLogP and total polar surface area (tPSA) (Fig. 2J).

Hotspot kinase profiling identified three analogs: UR241-2, PS-099, and PS-100, with ∼10-fold greater IRAK1/4 inhibition compared to JH-I-25 (Fig-2K). UR241-2 showed the highest potency and was selected for further study.

Kinome-wide profiling demonstrated limited off-target activity: 13 kinases were affected at 50 nM and 34 kinases at 500 nM (Fig-2L). In NanoBRET assays in HEK293 cells, IRAK4 was the most potently inhibited target (Fig-2M). UR241-2 inhibited IRAK1 with an IC₅₀ of 6.63 µM, while off-target kinases showed weaker inhibition, including FLT3 (5.07 µM), MAP4K1 (2.05 µM), MAP4K2 (114 nM), MAP4K3 (1.61 × 10⁻^5^ M), MAP4K5 (9.85 µM), and LRRK2 (283 nM).

Selectivity comparisons further supported this profile. UR241-2 was 67.9-fold more selective against FLT3 and 42-fold more selective against MAP4K1 relative to CTX0294885, and 33-fold more selective against MAP4K3. It showed 17,840-fold greater selectivity against MAP4K5 compared to Bosutinib and ∼16-fold greater selectivity against LRRK2 compared to CEP-701. Against IRAK1, UR241-2 exhibited 27-fold greater activity than pacritinib.

Together, kinome profiling and intracellular target engagement assays establish UR241-2 as a potent and highly selective IRAK4 inhibitor with minimal off-target activity.

### UR241-2 exhibits greater binding affinity and distinct interactions compared to JH-I-25

To elucidate the binding mode of UR241-2 and its analogs with IRAK4 and facilitate structure-activity relationship (SAR) development, we performed in silico docking simulations using UR241-2, PS-099, and PS-100 against the known IRAK4 crystal structure (Fig-3A). The interacting residues and nature of these interactions for each ligand are shown in Fig-3B, with JH-I-25 serving as a comparator. In these analyses, we examined aliphatic (green), aromatic (magenta), basic (blue), polar (sky blue), sulfur (yellow), hydrogen bond donor/acceptor (HBD/HBA) (dotted sky blue), hydrophobic (dotted green), and Van der Waals (VdW) (dotted cyan) interactions (Fig-3B, bottom). As expected from JH-I-25’s docking profile, the pyrazolo-pyridine core of all ligands occupied the same hinge region, though with modified residual interactions, highlighting a potential avenue for enhancing IRAK4 inhibition. Further analysis revealed that replacing the oxygen in JH-I-25’s morpholine with sulfur in thiomorpholine (PS-099) introduced a novel ASP114 hydrogen bond interaction, absent in JH-I-25. Although the pyrazole nitrogen atoms in JH-I-25 and PS-100 retained interactions with SER165, PHE34, ASN153, and ALA152, neither UR241-2 nor PS-099 engaged these residues, despite maintaining similar symmetry elements in morpholine, thiomorpholine, sulfoxide, and sulfone groups (Fig-3B). A superimposed view of all four ligands docked together is shown in Fig-3C.

Molecular dynamics simulations (1 ns) were performed to assess binding stability (Fig-3D-H). All ligands remained stably bound within the catalytic site, with ligand RMSD (root-mean-square deviation) values <2 Å, indicating minimal deviation from the docked pose (Fig-3D-E). Protein RMSD values similarly indicate structural stability of IRAK4 during simulation. RMSF (root-mean-square fluctuation) analysis showed limited residue-level fluctuation across all complexes (Fig-3F), and radius of gyration remained stable, consistent with compact ligand–protein complexes. Binding free energy calculations ranked UR241-2 as the highest-affinity ligand, followed by PS-100 and JH-I-25. PS-099 exhibited the lowest binding affinity, with a calculated energy of −10.5 kcal/mol (Fig-3H-I). Together, these data indicate that UR241-2 maintains stable binding within the IRAK4 catalytic pocket and achieves improved binding affinity relative to the parent scaffold through distinct interaction profiles.

**Figure-3:**
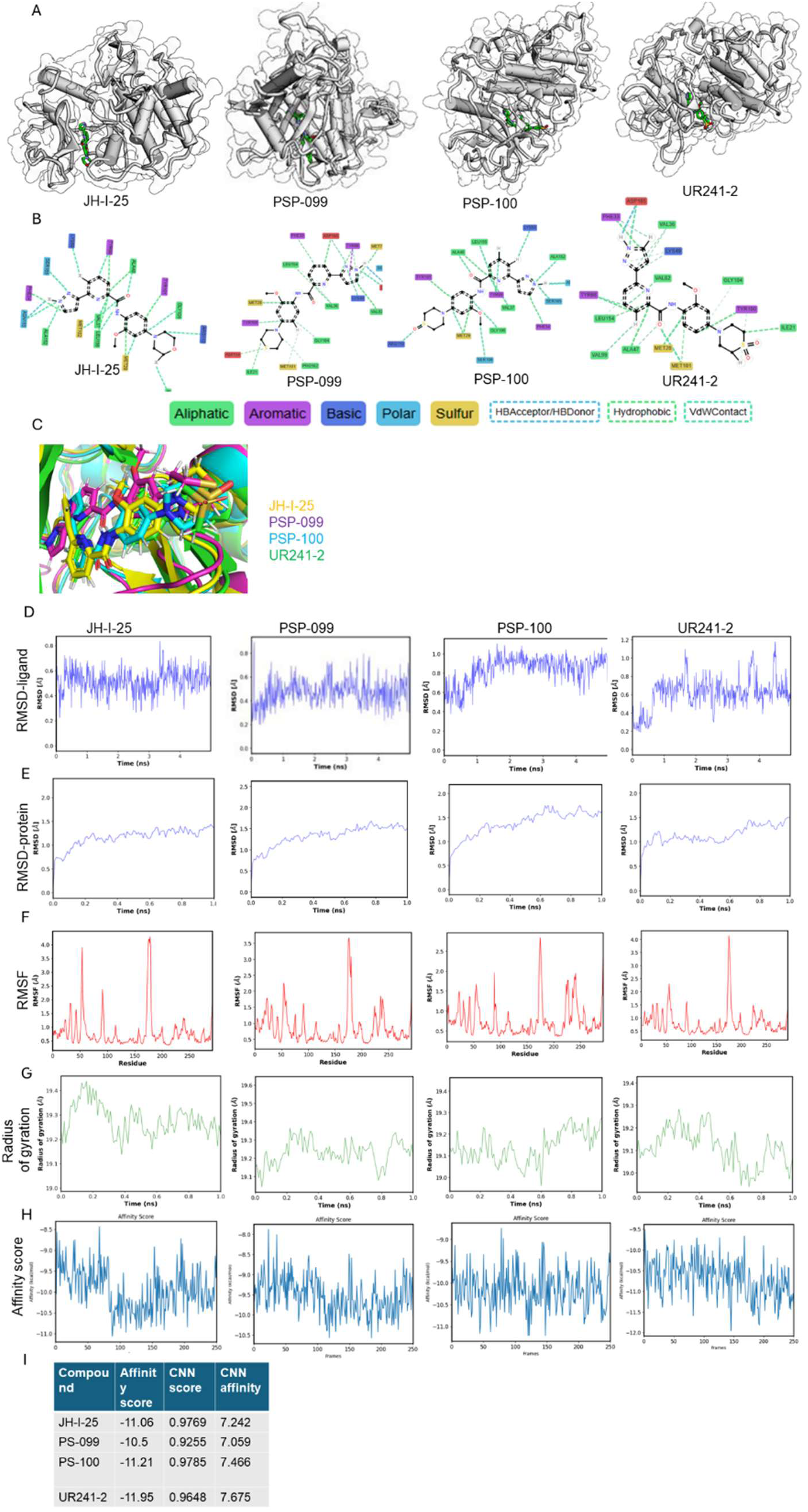
In silico docking and molecular simulations exhibit UR241-2’s interactions with IRAk4 protein. (**A**): JH-I-25 and its newer analogs UR241-2, PSP-099 and PSP-100 were docked individually to IRAK4 crystal structure using Gnina docking software, which is built on neural networks (CNNs) as a scoring function. (**B**): The interacting amino acid residues and their nature of interactions are shown. Green=aliphatic, magenta=aromatic, blue=basic, sky blue=polar, and yellow=sulfur. Red indicates unfavorable interactions. (**C**): JH-I-25, UR241-2, PSP-099 and PSP-100 analogs were docked together to IRAK4 protein using Gnina tools. (**D**): Root-mean square density (RMSD) simulations of JH-I-25, UR241-2, PSP-099 and PSP-100 ligands docked to IRAK4 falling within 2Å unit range are shown. RMSD-ligand is the root mean deviation of the bounded ligand compared to the ligand in the crystal structure. (**E**): Root-mean square density (RMSD) simulations of IRAK4 protein docked with JH-I-25, UR241-2, PSP-099 and PSP-100 ligands falling with 2Å unit range are shown. RMSD-Protein is the root mean deviation of the protein for each frame of the simulation compared to the crystal structure. Both **D** and **E** indicate that docking quality was acceptable. (**F**): RMSF (root mean square fluctuation) of JH-I-25, UR241-2, PSP-099 and PSP-100 ligands are shown. The ligands, when bound, induce the most notable structural changes in residues 172-181 of IRAK4 which are reflected in the spike in the RMSF charts. RMSF is the mean fluctuation of IRAK4 protein residues after binding of ligands. RMSF indicates the conformational flexibility of the complex. The time/frame element is removed from RMSF in order to show an average fluctuation of the residues. (**H**): The affinity score measures the affinity of a ligand (in that particular frame) binding to the protein. The frames represent different conformations that a ligand may adopt. The affinity is measured in kCal/mol and the lowest frame (lowest energy) is selected as the most stable binder. UR241-2 showed the best affinity score (-11.95), hence was considered the most stable binder ligand of IRAK4 crystal structure among the compounds tested.

### UR241-2 inhibits IL-1β–induced IRAK4 phosphorylation in EOC cells

We first assessed the effect of UR241-2 on IL-1β–induced IRAK4 activation in EOC cell-lines. In murine EOC cell lines (HGS-1 and HGS-3), pre-treatment with UR241-2 (20–150 nM, 4 h) inhibited phosphorylation of IRAK4 following stimulation with mIL-1β (10 ng) (Fig-4A). In human HCH-1 EOC cells, UR241-2 (5 and 15 nM, 4 h) similarly suppressed IRAK4 phosphorylation induced by hIL-1β (10 ng, 45 min) (Fig-4B).

**Figure-4:**
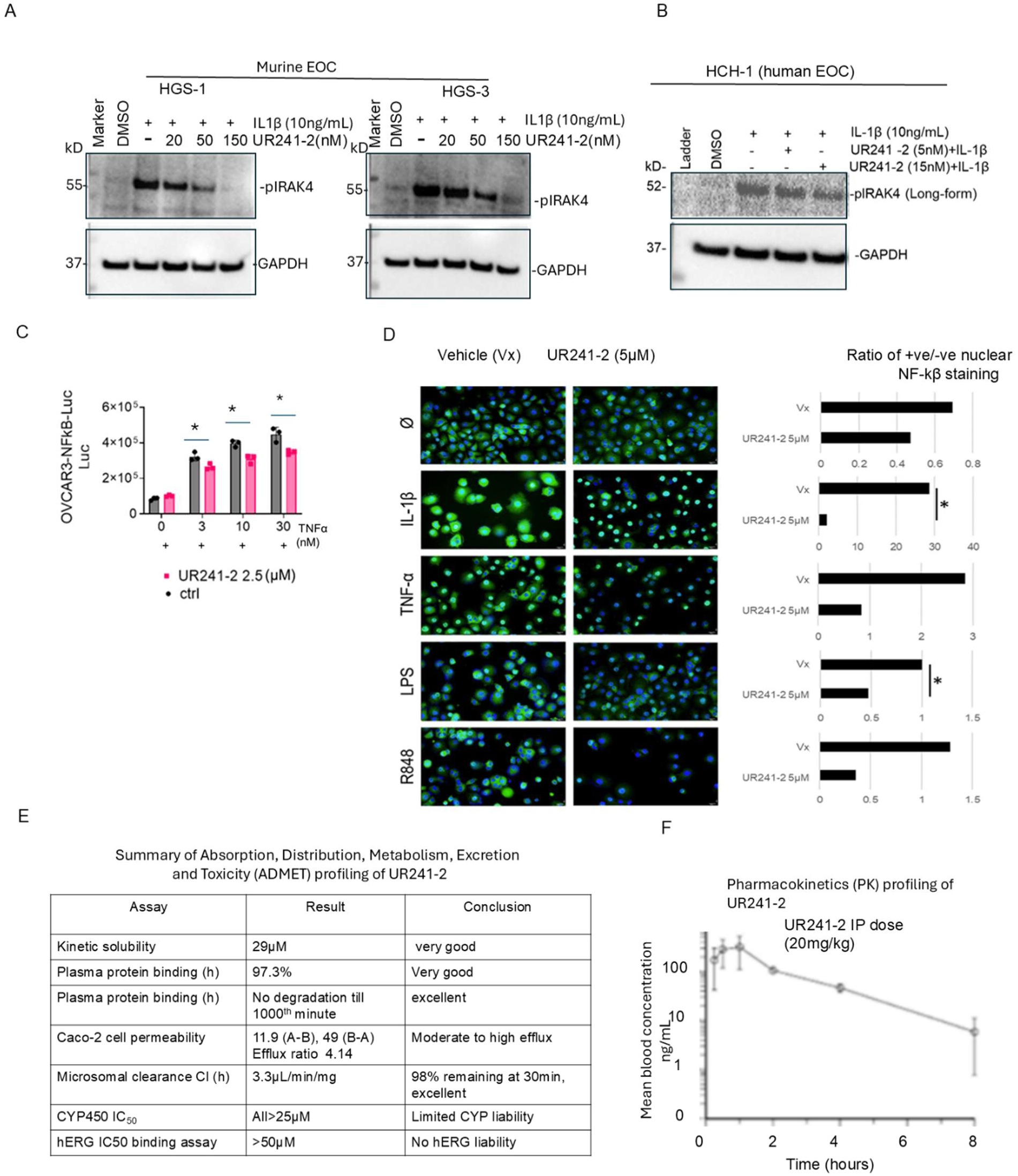
UR241-2 blocks IL1β induced IRAK4 phosphorylation. (**A-B**): UR241-2 (5-150nM) treatment for 4hrs blocks human and murine IL1β (5nM, 30 minutes) induced IRAK4-phosphorylation in both human (HCH-1) and murine high-grade serous EOC cells (HGS-1 and -3) cells. (**C**): UR241-2 (2.5μM) treatment reduced NF-κβ luciferase reporter activity induced by human TNF-α (3, 10 and 30nM) in stably transfected OVCAR-3-NF-κβ-Luc cell-lines. * indicates p=0.01 to 0.008 (two tailed unpaired T-test). (**J**): Similarly, UR241-2 (2.5μM) treatment, IL-1β (5ng), TNF-α (10ng), LPS (100nM) and R848 (TLR-7/8 agonist, 10μM). Ratio of positive/negative NF-κβ nuclei are shown. NF-κβ (Green) nuclear migration was inhibited significantly in IL-1β and LPS stimulated cells. The number of NF-κβ nuclear positive cells were counted using ImageJ software. * indicates <0.05 (Student T-test). (**D**): Summary of ADMET profiling of UR241-2. ADMET characteristics of UR241-2 were examined in the laboratories of Curia Inc, NY, USA. Complete ADMET along with positive controls is shown in the Supplementary Figure-3. (**E**): Pharmacokinetic PK profiling of UR241-2 in CD-1 mice. Mice were intraperitoneally injected with UR241-2 and blood concentration of UR241-2 was quantified by HPLC at the indicated hours. See also Supplementary Figure-3.

### UR241-2 suppresses NF-κβ activation downstream of IL-1/TLR signaling

We next examined NF-κβ signaling, a key downstream effector of the IL-1/TLR/MYD88/IRAK1/4 pathway. In OVCAR-3 cells stably expressing an NF-κβ luciferase reporter^46^, UR241-2 (2.5 µM) reduced NF-κβ activity following TNF-α stimulation at 3, 10, and 30 nM (Fig-4C). To assess nuclear translocation, OVCAR-3 cells were treated with UR241-2 (5 µM) and stimulated with IL-1β, TNF-α, LPS, or the TLR7/8 agonist R848. UR241-2 significantly reduced nuclear NF-κβ levels compared to vehicle controls under all conditions (Fig-4D). Specifically, the nuclear NF-κβ positive/negative expression ratio decreased following IL-1β and LPS stimulation and similarly reduced in response to TNF-α and R848.

### UR241-2 exhibits favorable ADMET properties

ADMET profiling of UR241-2 is summarized in Fig-4E, with detailed data in Supplementary Fig-3. At pH 7.4, UR241-2 showed high kinetic solubility (29 µM) in PBS containing 1% DMSO (Supplementary Fig-3A). In microsomal stability assays, UR241-2 was stable in human liver microsomes (t₁/₂ = 208.9 min; CLint = 3.33 µL/min/mg protein) but showed faster clearance in mouse liver microsomes (t₁/₂ = 8.70 min; CLint = 79.7 µL/min/mg protein) (Supplementary Fig-3B).

CYP inhibition assays across eight isoforms (1A2, 2B6, 2C8, 2C9, 2C19, 2D6, 3A4M, 3A4T) showed weak inhibition, with IC₅₀ values ranging from 30 to >100 µM (Supplementary Fig-3C). Plasma protein binding was high in both human (97.4 ± 0.3%) and mice (93.8 ± 0.8%) plasma (Supplementary Fig-3D).

In CaCo-2 assays, UR241-2 showed an efflux ratio of 4.14, consistent with classification as an efflux substrate (Supplementary Fig-3E). hERG channel assay showed no significant inhibition up to 100 µM (EC₅₀ = 1.01 × 10^-4^), whereas the positive control E-4031 showed strong inhibition (EC₅₀ = 1.82 × 10^-8^) (Supplementary Fig-3F).

### UR241-2 demonstrates acceptable plasma levels in a mouse pharmacokinetic (PK) study

To assess serum levels over time, we conducted a pilot PK study in CD-1 mice. UR241-2 was administered at 20 mg/kg intraperitoneally (IP) and 1 mg/kg intravenously (IV). Plasma concentrations were analyzed using LC-MS. As shown in Supplementary Fig-3G, IV administration resulted in rapid clearance, whereas IP administration (20 mg/kg) maintained plasma levels above 8 ng/mL for up to 8 hours (Fig-4F).

### UR241-2 suppresses EOC growth in vitro and in vivo without overt toxicity

We first assessed the effect of UR241-2 on colony formation in EOC cell lines. In SKOV-3 cells, colony size was reduced at 5 µM (p = 0.0131), 10 µM (p = 0.0437), and 20 µM (p = 0.0077) (Fig-5A, upper). In OVCAR-3 cells, colony size was reduced at 5 µM (p = 0.0488), with greater suppression at 10 µM (p < 0.0001 vs. control; p = 0.0142 vs. 5 µM) and 20 µM (p < 0.0001 vs. control; p = 0.0104 vs. 5 µM) (Fig-5A, middle). In OVCAR-8 cells, colony size was reduced at 5 µM (p = 0.0038), with further reductions at 10 µM (p = 0.0012 vs. control; p = 0.0422 vs. 5 µM) and 20 µM (p = 0.0055 vs. control; p = 0.0116 vs. 5 µM) (Fig-5A, lower). Average colony size across conditions is summarized in Fig-5B.

Cell proliferation was assessed using SRB assay^47^. Treatment with UR241-2 (20–60 µM, 48 h) resulted in a dose-dependent reduction in viability across ES2, HCH1, SKOV-3, OVCAR-3, and OVCAR-8 cell lines (Fig-5C). To assess cell division, we measured Histone-H3 phosphorylation. UR241-2 reduced the proportion of mitotic cells in OVCAR-3 (p = 0.0024 at 20 µM) and SKOV-3 cells (p = 0.0047 at 10 µM; p = 0.0003 at 20 µM) (Fig-5D).

**Figure-5:**
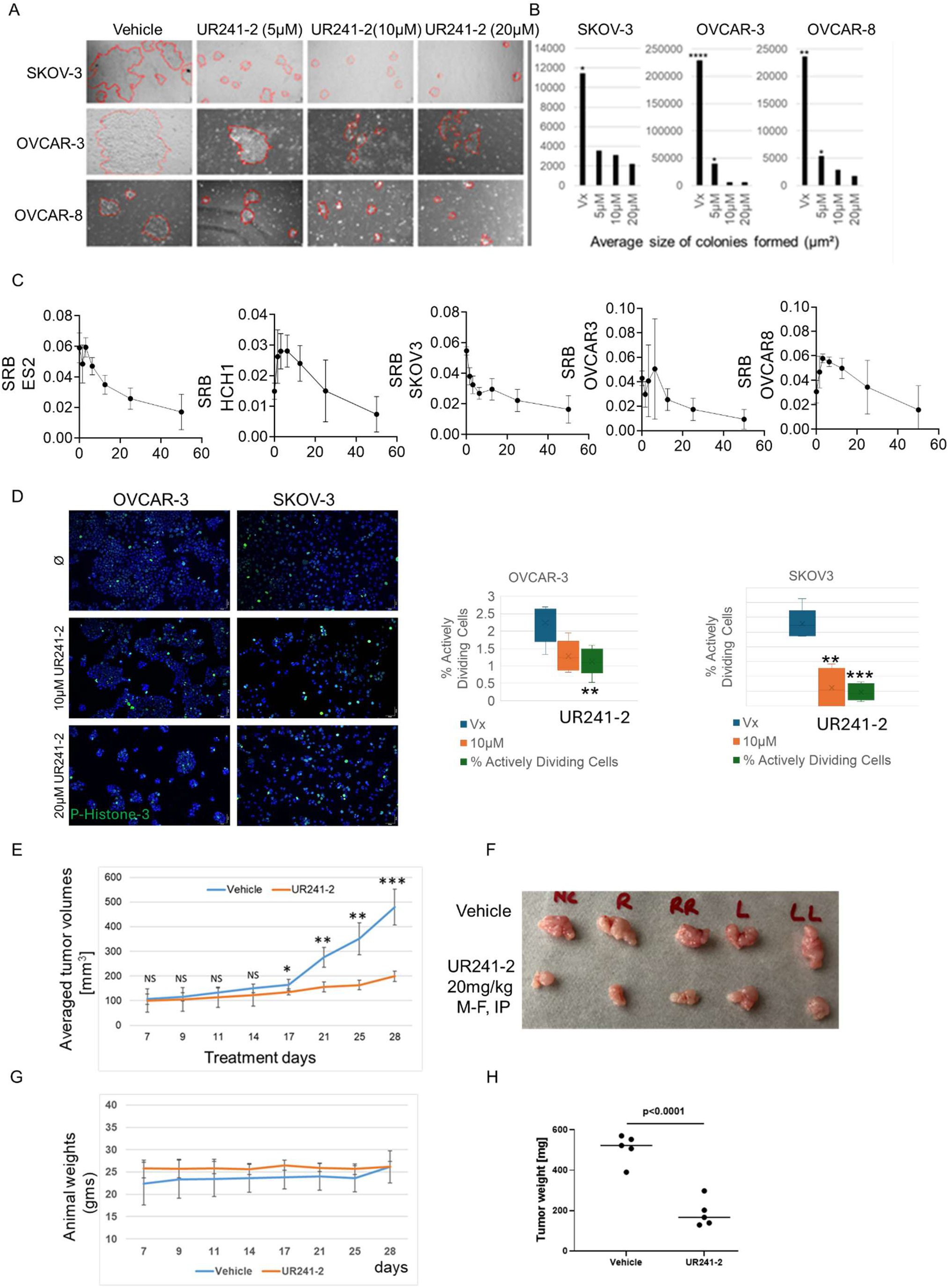
UR241-2 treatment colonies, proliferation, mitotic index *in vitro* and tumor growth *in vivo*. (**A**): UR241-2 (5, 10 and 20 µM) treatment for 7 days blocks colonies formed from 2000 cells seeded in a 6 well plate compared to control. One representative image of multiple images of each well that were collected are shown. (**B**): Average size of colonies in vehicle versus 5, 10 and 20 μM groups were analyzed by ImageJ software. Non-parametric T-test was conducted to determine the statistical differences between the control and various treatment groups. * indicates <0.05 (Student T-test). (**C**): UR241-2 (20-60μM) treatment reduced cell viability in ES2, HCH1, SKOV-3, OVCAR-8 and OVCAR-3 EOC cells. The cell viability was measured by SRB assay which measures the total protein synthesis in drug treated cells compared to control vehicle. (**D**): UR241-2 (10-20μM) treatment reduced cell division in SKOV-3 and OVCAR-3 cells. The dividing cells were captured by pHistone-3 staining. SKOV-3 and OVCAR-3 cells (10,000/well) were seeded in an 8-well EasyCell chamber slides overnight. The cells were treated with vehicle and UR241-2 (10-20μM). Cells were fixed after 24-hours using neutral buffered formalin solution (35µL) for 20 minutes. The media was removed, and cells were washed repeatedly with TBST (500µL/5 min). Fixed cells were stained with p-Histone-H3 antibody (Cell Signaling Technology, cat#9701, 1:500 dilution, overnight) and then counterstained with Dylight-488(Vector laboratories, cat#DI-1488) secondary antibody (1:2000). The cells were washed repeatedly with TBST (500µL/5 min). Washed cells were stained again with the Vectashield mounting media containing DAPI (Vector Lab, cat#H-1200-10), protected with a cover glass slide, then imaged on an Olympus BX41 microscope. All images taken with a 10x ocular and 20x objective. All cells in each field were manually counted and assessed as either actively dividing (metaphase, anaphase, telophase) or not. OVCAR-3 vehicle group had significantly more actively dividing cells than the 20μM treatment group (p=0.0024). SKOV3 vehicle group had significantly more actively dividing cells than both 10μM and 20μM treatment groups (**p=0.0047, ***p=0.0003). (**E**): UR241-2 treatment (20mg, M-F, IP) reduced the growth of SKOV-3 cells derived xenografts growing in NSG mice. 10 NSG mice implanted subcutaneously with SKOV-3 cells (1 million/mice in DMEM+Matrigel (1:1, 100µL/mice) were randomized and treated for 28-days. Tumor sizes were measured on the days indicated in the X-axis. Mice were euthanized and tumors and peripheral blood were harvested. Averaged tumor volumes differed significantly between the control and treatment groups. *=p<0.05, **=p<0.005, ***=p<0.0005. (**F**): Images of the tumors formed in vehicle and UR241-2 treated mice are shown. (**G**): Animal weights did not differ. (**H**): Tumor weights between vehicle and treated groups differed significantly. The tumor weights of the individual mice in the control and treatment groups were plotted using GraphPrism Version 8.0. *p*=0.0001.

We next evaluated in-vivo efficacy using a SKOV-3 xenograft model. Based on pharmacokinetic data (Supplementary Fig-3G), UR241-2 was administered at 20 mg/kg (intraperitoneal, once daily, Monday–Friday). Tumor growth was reduced beginning on Day 17 (p < 0.05), with significantly smaller tumors by Day 28 compared to vehicle-treated controls (p < 0.0005) (Fig-5E). Final tumor weights were also reduced (p < 0.0001) (Fig-5F, H). UR241-2 treatment did not affect body weight or overall well-being (Fig-5G). Hemoglobin levels in peripheral blood were unchanged between treated and control groups (Supplementary Fig-4)

### UR241-2 reduces injury-associated tumor growth and modulates inflammatory immune populations

We evaluated UR241-2 in a syngeneic MIM model of EOC that recapitulates tumor adhesion at sites of injury. Intraperitoneal implantation of serous murine ovarian cancer HGS-3 cells^48^ mimics fallopian tube-derived EOC^49^. UR241-2 treatment reduced tumor size at the injury site (Fig-6A) but did not reduce omental tumor burden (Fig-6B), indicating that injury- or inflammation-associated signaling contributes to IRAK4-dependent tumor growth.

To assess immune modulation, leukocytes from peritoneal lavage and tumor tissue of HGS-3 tumor-bearing mice were analyzed by flow cytometry (Fig-6C). Representative plots of F4/80 and MHCII-expressing myeloid cells are shown for vehicle- and UR241-2-treated mice. UR241-2 increased both the percentage and number of MHCII^hi^ macrophages in the peritoneal lavage (Fig-6D, left), while reducing MHCII^low^ macrophages (Fig-6D, right), consistent with a shift toward antigen-presenting myeloid populations. UR241-2 also reduced the proportion of F4/80^hi^CD206^+^-M2-like macrophages in the peritoneal cavity (Fig-6E).

**Figure-6:**
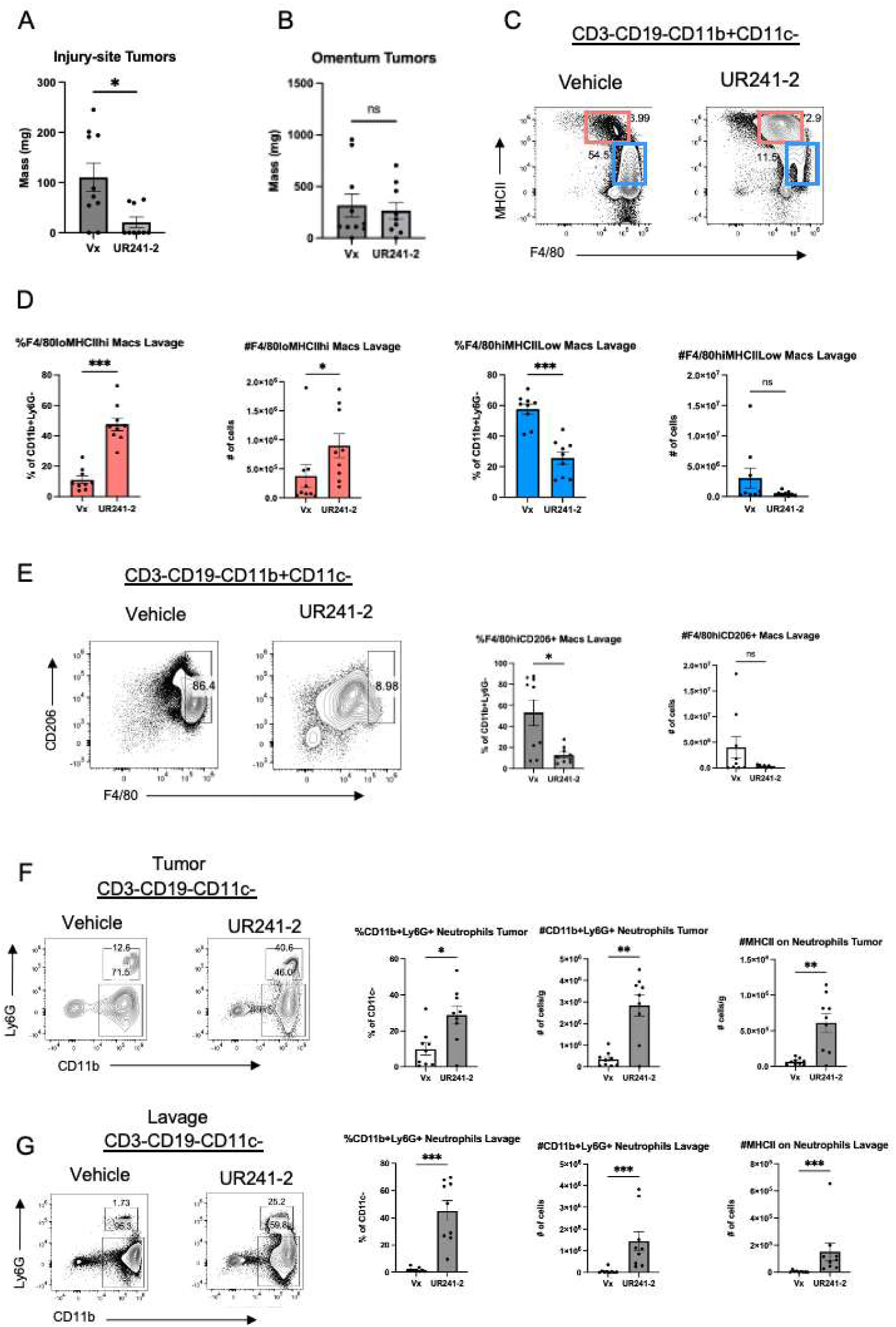
IRAK4 inhibition via UR241-2 decreases tumor burden at the needle injury site and increases the number of MHCII expressing myeloid cells in the peritoneal cavity and tumor. (**A**): Tumor weights on the needle injury site were significantly lower in the UR241-2 treatment group than vehicle treated animals. Statistical significance was determined using Mann-Whitney test, *p < 0.033, **p < 0.002, ***p < 0.001. (**B**): Tumor weights on omentum did not differ between UR241-2 treatment group than vehicle treated animals (NS). Two of the three replicates are combined and shown. IRAK4 inhibition via UR241-2 increases the number of MHCII expressing myeloid cells and decreases MHCII low and CD206+ macrophages in the peritoneal cavity. Leukocytes were isolated from the peritoneal lavage or tumor of HGS-3-tumor bearing mice and stained for flow cytometry. (**C**) Representative flow cytometry plots of F4/80^lo^MHCII^hi^ M1 macrophages and F4/80^hi^MHCII^Low^ macrophages from the peritoneal lavage of HGS-3 tumor bearing mice treated with Vehicle or UR241-2. (**D**) Percentage and cell number of F4/80^lo^MHCII^hi^ M1 macrophages and F4/80^hi^MHCII^Low^ macrophages from C. **(E)** Representative flow cytometry plots of F4/80^hi^CD206^+^ macrophages from the peritoneal lavage of HGS-3 tumor bearing mice treated with Vehicle (Vx) or UR241-2. (**E-right**) Percentage and cell number of F4/80^hi^CD206^+^ macrophages. IRAK4 inhibition reduces neutrophil numbers in the peritoneal cavity and tumors of HGS3 bearing mice: **(F**): Flow cytometry plots showing CD11b^+^Ly6G^+^ neutrophils and MHCII^+^ neutrophils in the HGS-3 tumor. Percentage and cell number per gram are quantified to the right. (**G**): Flow cytometry plots showing CD11b^+^Ly6G^+^ and MHCII^+^ neutrophils in the peritoneal lavage of HGS-3 tumor bearing mice. Percentage and cell number are quantified to the right. N= 9 mice per group from two independent experiments. Gating strategies are shown in Supplementary Data-7. One mouse from the UR241-2 group died prior to analysis.

Notably, UR241-2 increased neutrophil populations in both tumors and peritoneal lavage (Fig-6F, G). In addition, MHCII expression on neutrophils was higher in UR241-2-treated mice in both compartments, resulting in an increased number of MHCII^+^ neutrophils (Fig-6F, G). Given the functional plasticity of neutrophils, including their ability to present antigens on MHCII^50^, this increase is consistent with a shift toward a more inflammatory, anti-tumor immune state.

Together, these data indicate that while IL-1R1 signaling contributes to inflammatory responses in the peritoneal and tumor microenvironment, IRAK4 inhibition by UR241-2 reshapes this response by increasing MHCII-expressing cells and reducing CD206^+^ macrophages.

### UR241-2 alters ECM and neutrophil-related gene expression in tumors

Bulk RNA-sequencing of UR241-2-treated tumors revealed significant changes in gene expression linked to extracellular matrix (ECM) organization and neutrophil-mediated immunity. Tumors from Fig-6A were analyzed, and principal component analysis (PCA) showed clear separation between vehicle- and UR241-2-treated samples (Fig-7A). Volcano plots (Fig-7B, 7C) highlighted differentially expressed genes. Among the top 30 downregulated pathways (adjusted p-values), ECM organization genes-critical for tumor cell adhesion, were strongly suppressed (Fig-7D). In contrast, neutrophil-associated pathways, including neutrophil-mediated immunity and activation, were significantly upregulated (Fig-7E). Heatmaps confirmed global transcriptional changes (Fig-7F) and ECM gene suppression in treated tumors (Fig-7G). Functionally, UR241-2 reduced HGS-3 adhesion to a collagen-coated basement membrane in a dose-dependent manner (Fig-7H). Cell migration was also significantly inhibited at concentrations as low as 250 nM (Fig-7I). These results align with the flow cytometry data, showing increased neutrophil populations and MHCII^+^ cells in-vivo. Together, this data indicates that UR241-2 downregulates ECM-related genes, impairing tumor cell adhesion and invasion, while simultaneously upregulating neutrophil activation pathways, promoting pro-inflammatory, anti-tumor immune responses in the peritoneal cavity.

**Figure-7:**
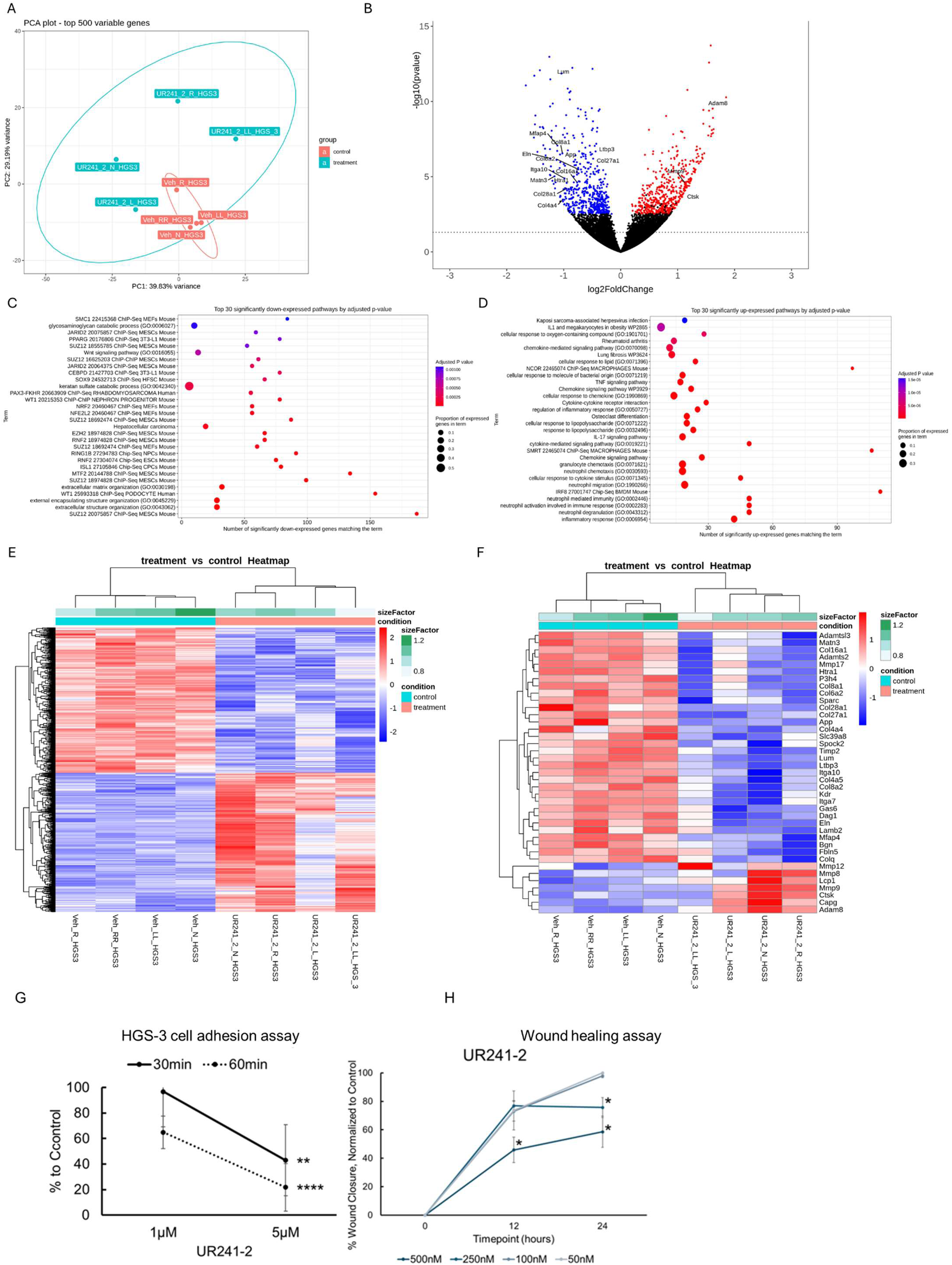
Bulk-seq of UR241-2 treated tumors show downregulation of ECM genes and upregulation of neutrophil activation genes. **(A):** PCA plot of Bulk-Seq analysis of control and UR241-2 treated tumors. (**B**): Volcano plot of genes, overexpressed and downregulated between control and treatment groups are shown. Downregulated genes belonging to ECM family are shown. (**C**): Gene-families downregulated by UR241-2 are shown. (**D**): Gene-families upregulated by UR241-2 are shown. (**E**): Heatmap of the genes altered between control and treatment. (**F**): Heatmap of the ECM family of genes downregulated by UR241-2 are shown. (**G**): HGS-3 cells treated with UR241-2 demonstrated an inhibited ability to adhere to collagen compared to control, with cells treated at 5μM showing significantly less adhesion than the 1μM group at both the sixty (p<0.00005) and thirty (p=0.00068) minute timepoints. (**H**): Cell migration in HGS-3 cells treated with 500nM UR241-2 were significantly inhibited compared to control in just 12 hours (p<0.02), and both the 250nM and 500nM groups showed a significantly greater impact than the 100nM, 50nM, and control treatment groups at 24 hours (p<0.05).

## Discussion

While the clinical phenomenon of tumor cells seeding at injured/inflamed sites is observed across multiple cancers, among sizeable patient populations and predicts residual, aggressive and recurrent diseases, it remains poorly understudied and lacks targeted therapies. In ovarian cancer, this clinical event has been lacking in-depth mechanistic and therapeutic advances, being managed by radiation, chemotherapy, or surgical resection. Our study demonstrated the roles of IL-1β/IL1R1/IRAK4 signaling in EOC seeding^51–52^ and tumor growth at mechanically injured sites that recreate the inflammatory milieu of the peritoneum during metastasis^53^.

These studies were enabled by the development of a needle-injury–induced MIM model, a reproducible, non-surgical syngeneic platform that recapitulates cell adhesion, ECM remodeling, and immune microenvironmental changes likely occurring during metastasis. The MIM model mimics port site metastasis (PSM) observed in gynecologic and other surgeries and allows temporal analysis of ECM-immune-tumor interactions. Compared with the burn-induced ovarian surgical model described by Jia et al.^1^, the MIM model achieves comparable biological relevance while limiting procedures to a brief, momentary needle injury instead of survival surgeries using wounded or burned ovaries. Using this platform, we demonstrate that UR241-2, a highly selective IRAK4 inhibitor, prevents and mitigates tumor seeding at sites of inflammation. This model can now be utilized for identification of novel therapies to block or interfere with inflammation driven metastatic seeding across most other malignancies.

Theoretically, IL-1β/IL-1R1/IRAK4 drives EOC seeding, agents such as Anakinra (recombinant IL-1 receptor antagonist), Canakinumab (IL-1β neutralizing antibody), or UR241-2 each could prevent seeding driven by inflammation. However, UR241-2 robustly reprograms the peritoneal immune milieu, providing durable anti-tumor activity similar to Il1r1-deficient mice (Fig-1N). Notably, UR241-2 induces MHCII expression on neutrophils, enabling antigen-presenting cell (APC) functionality without exogenous cytokines such as IFN-γ or GM-CSF. Bulk RNA-seq confirmed activation of neutrophil antigen-presentation programs (Fig-7). However, additional functional assays will be needed to confirm antigen-presenting capacity, but this ability to induce immune-reprogramming positions UR241-2 as a promising candidate for therapy combinations with immune checkpoint inhibitors (ICIs), such as pembrolizumab, potentially enabling neutrophils to traffic to lymphoid tissues, prevent metastasis^54^, and amplify anti-tumor immune responses including taking the advantages of combination with NF-kβ tareting therapies (Model-1).

UR241-2 exhibits favorable pharmacologic and ADMET properties. It inhibits IL-1β-induced IRAK4 phosphorylation, selectively suppresses the long-form IRAK4 without affecting the short isoform, blocks NF-κβ signaling, and reduces adhesion, proliferation, and division of EOC cells, with significant effects on colony formation and SKOV-3 xenograft growth. Importantly, UR241-2 treatment did not cause body weight loss or hematologic toxicity. Pharmacokinetic data indicates that 20 mg/kg IP administration maintains ∼8 pM plasma levels for 8 hours in C57BL/6 mice (Supplementary Fig-3G), correlating plasma exposure with in-vivo efficacy.

While UR241-2 demonstrates promising multi-level therapeutic activity, tumors did not completely regress. Therefore, its clinical utility may lie primarily in combination with chemotherapy or immunotherapy, particularly after the first round of surgery and systemic therapy. Combination approaches generally produce stronger and more durable responses in malignancies, including EOC. For example, Anakinra enhances fluorouracil sensitivity in KRAS-mutant colon cancer^55^ and has been tested as part of the AGAP regimen (gemcitabine, nab-paclitaxel, cisplatin, Anakinra) in localized pancreatic ductal adenocarcinoma^56^. Similarly, CA4948, a non-selective IRAK4 inhibitor, is under clinical evaluation with PD-L1/PD-1 blockade in pancreatic, esophageal, gastric, bladder cancers and AML. For EOC, future studies will examine UR241-2 in combination with paclitaxel, the first-line standard of care. Paclitaxel activates TLR4, promoting chemoresistance, and we speculate that paclitaxel+UR241-2 may block TLR4-mediated resistance, enhancing chemotherapy efficacy and overall survival.

A limitation of this study is its singular focus on EOC adhesion to injured mesothelium, without directly addressing IL1/IL1R1 roles in omental seeding, as the omentum was uninjured in the MIM model. Consistently, neither Il1/Il1r1 loss-of-function nor UR241-2 reduced omental tumor burden, suggesting that adhesion at this site may be less dependent on IL-1β/IL1R1/IRAK4 and injury-associated inflammation, although tissue- or context-dependent roles cannot be excluded. Notably, omental macrophages are known to support EOC migration and colonization^57^. While UR241-2 altered peritoneal macrophage surface markers, functional impact on omental macrophage polarization was not assessed and remains for future study. Importantly, this lack of effect on omental tumor burden does not diminish the translational relevance of UR241-2 or other IRAK4 inhibitors in EOC, as the omentum is typically removed during cytoreductive surgery. This leaves the peritoneum as a common site of recurrence, underscoring the need for IRAK4-targeted therapies to prevent metastatic seeding post-surgery. The ability of UR241-2 to reduce peritoneal metastasis, a major driver of mortality in post-surgical recurrent EOC, highlights its potential clinical value.

## Methods

### Microarray database analysis

Kaplan-Meier survival outcomes among EOC patients (Fig-1A, 1B, 1Q and 1R) were analyzed using the databases (tcga-381-tpm-gencode36) and tools available at R2-Genomics and Visualization Platform (https://hgserver1.amc.nl/) using the system selected gene expression cut-off points. For determining the expression of IRAK4 the database (Mori-171-rma_sketch_hugene10t) was also analyzed. Gent2 database (http://gent2.appex.kr/gent2/) was used to compare the IRAK4 (Fig-1O) differences between normal ovaries versus malignant ovaries. Gent2 database was also used to compare the IRAK4 expression differences in EOC stages (date accessed: 10/2/2023) (Fig-1P). IRAK4 mRNA correlation with E-cadherin mRNA was determined using R2-Genomics and Visualization Platform (https://hgserver1.amc.nl/) using the system selected gene expression cut-off points (Fig-2F). Cut-off points were not altered.

### MiM model

3-7 million HGS-3 murine high-grade serous EOC cells suspended in serum and antibiotic free DMEM were injected intraperitoneally in C57/BL6 mice or their derivative Il1r1^KO^, Il1rn^KO^ and Nlrp3^KO^ mice using 21G needles. Mice are monitored for 40-60 days when most of the mice will present tumor nodules at the site of needle injury. Skin of the mice was regularly shaved carefully to remove the fur until the tumors became palpable and measurable.

### EOC tumor burden in syngeneic Il1r1^KO^, Il1rn^KO^ and Nlrp3^KO^ mice

*Il1r1*^KO^ (B6.129S7-Il1r1tm1Imx/J), *Il1rn*^KO^ (Fig-1E) (B6.129S-Il1rn tm1Dih/J) (Fig-1K) and *Nlrp3*^KO^ (Strain #:021302 - B6.129S6-Nlrp3tm1Bhk/J) mice were bred in-house from the breeders obtained from Jax laboratories. Homozygous *Il1r1*^KO^ (n=5), *Il1rn*^KO^ (N=5) and *Nlrp3*^KO^ (n=5) mice were implanted with HGS-3 mouse HGS-3 EOC cells (3-4.5 million/mice) intraperitoneally. Similarly, WT C57BL/6 mice (n=5) were also injected intraperitoneally with HGS-3 murine HGS EOC cells (3-4.5 million/mice) using 21G sterile needles. After 45 days, mice were bled via mandibular vein and ∼25µL of blood collected for complete blood counts (CBC) was analyzed in EDTA-tubes using Element-HTS+ system. Mice were euthanized soon after and photographed to record the tumors formed at the injury sites. Spleen, omentum, and peritoneal lavage were collected from each mouse. Omentum weights were recorded and weighed. Spleen was crushed in serum-free RPMI1640 medium (Gibco, cat#: 22400-089) and filtered. Tumors/omentum were processed similarly. Harvested cells were broken down into single cell suspension and analyzed by flow cytometry following the conditions described below.

### Flow cytometry

Omentum and tumors were minced in plain RPMI 1640 media and incubated in collagenase (2mg/ml), DNase (25ug/ml), and hyaluronidase (0.1mg/ml) for 30min-1hr at 37^0^C. The suspension was strained through 70µm strainers, diluted with 10ml of cold complete RPMI (10% FBS) and centrifuged at 300 x g for 5 mins at 4^0^C. The cells were resuspended in PBS for cell counting and stained for flow cytometry. The spleen was isolated and crushed through a 70µm strainer in cold complete RPMI (10% FBS) and transferred to tubes and the cells were centrifuged at 300 x g for 5 mins at 4^0^C. The resulting pellet was resuspended in 1X Red blood cell lysis buffer for 4-5mins at room temperature with occasional shaking. The suspension was diluted with complete RPMI (10% FBS) and centrifuged at 300 x g for 5 mins at 4^0^C. The pellet was resuspended in RPMI (10% FBS) for cell counting and stained for flow cytometry. The peritoneal lavage was collected using a 5 mL syringe and 20G needle to inject 5 mL of PBS into the peritoneal cavity. The abdomen was massaged for 10-15 seconds, and the peritoneal lavage was collected using the needle. The needle was removed, and the resulting suspension dispersed into a tube on ice. The suspension was centrifuged at 300 x g for 5 mins at 4°C and resuspended in 10 mL of PBS and passed through a 70 µm strainer. The suspension was centrifuged at 300 x g for 5 mins at 4^0^C and resuspended in PBS for cell counting and stained for flow cytometry. After staining, all tissues were fixed and analyzed using a Cytek Aurora Full Spectrum Flow Cytometer. The staining antibodies were used at a dilution of 1:200. The list of antibodies, catalog numbers and sources are described in Supplementary List-5.

### Characterization data of UR241-2 and analogs

The synthesis of UR241-2 and analogs (PSP-099 and PSP-100) (Fig-2I) was achieved in milli gram scale. The compounds were characterized by ^1^H- and ^13^C-NMR and Mass spectrometry (MS) (Supplementary data-6). Purity was established by HPLC. The compounds tested in this study were each more than 97% purity by HPLC.

### Hotspot Kinase, Global kinome analysis and cell based nanoBret kinase inhibition assay

UR241-2 and analogs were screened for the IRAK1, -2, and -4 kinase activity in 10-dose singlet curve mode using the Hotspot kinase assay offered by Reaction Biology Inc. (Fig-2K). Selectivity of UR241-2 was analyzed against more than 680 kinases in a Global kinome screening assay offered by Reaction Biology Inc. at 3 escalated doses. Dendrograms of kinase activities at 50 and 500nM dose of UR241-2 were prepared using the KinMap^58^ (http://www.kinhub.org/kinmap/) (Fig-2L). The specificity of UR241-2 against IRAK1, IRAK4, LRRK2, FLT3, MAP4K1, MAP4K2, MAP4K3, MAP4K5. The kinases that showed comparable kinase inhibition in Hotspot kinase screening *in vitro* were determined using HEK293-cell-based nanoBret assay (Fig-2M).

### In silico docking and molecular docking simulations

The crystal structure of IRAK4 was downloaded as PDB file from the PDB bank (RCSB ID: 2NRU). The protein was prepared prior to docking, which involved selecting a single chain out of the four identical ones present. Further preparation was handled using the GNINA docking suite^59^ (https://github.com/gnina) with machine learning (ML) hosted on Google Colab. Next, the docking grid box was created to serve as the search area for the GNINA docking suite. The T12 ligand was selected as the center of the grid box. This area is the ATP binding site and was chosen due to the design of the molecule serving as a competitive inhibitor, completing protein preparation. Ligand preparation began conversion from SMILES to SDF. The SDF file was uploaded to Google Drive. Further preparation of the ligand continued with the protonation of the ligand and the subsequent geometry optimization using Torch ANI, with the convergence threshold set to 0.0001, thus completing the ligand preparation. Next, we selected our docking parameters for GNINA. Exhaustiveness was set to 64, buffer space was set to 4 angstroms, and refinement scoring was used. Docking was then performed via the GNINA docking suite and obtained multiple poses. The best pose was selected based on the highest affinity and the predicted location of our ligand (Fig: 3A-C). Ligands outside of the ATP binding domain were disregarded. This completed the docking phase. The docked ligand and protein were then moved to undergo simulations. Simulations were done using the OpenMM engine. A notebook made by the “Making it rain” team provided written code for the engine on Google Colab. Parameters for topology generation were set as following: the NaCl concentration was set 0.15 mol, padding distance was set to 4, the force field used was the AMBER99SB, the water type model was set to SPC/E, pH was set to 7.4, and finally gaff-2.11 was used to generate ligand force field. MD equilibration was done prior to the MD simulation. Both were simulated at a temperature of 298K and 1-bar pressure. The equilibration process was conducted for 0.5 nanoseconds, and the simulation was conducted for 5 nanoseconds. Images were generated by the libraries within the notebook (Fig:3D-I). Physicochemical properties of the compounds were calculated with ChemDraw software using the chemical analysis tools (Fig-2J).

### Effects of UR241-2 on NF-κB activity in EOC cells

OVCAR-3 cells (20,000/well) were seeded in 8-well chambers slides and cultured in DMEM (Gibco, cat#: 11885-084) supplemented with 10% FBS (Avantor, cat# 89510-186), 1% penicillin/streptomycin (Gibco, cat#:15140-122). Media was replaced with serum free DMEM media cells were stimulated with either IL1β (5ng), TNF-α (5ng), LPS (5µg), R848 (10µM), or negative control, for 24 hours prior to treatment start. Each group was either treated with 5μM UR241-2 or DMSO vehicle for 24 hours prior to analysis. Cells were then fixed with neutral formalin for 25 minutes at 4°C and washed with PBST(3x500µL). The cells were stained with NF-κβ p65 (D14E12) XP® Rabbit mAb primary antibody (Cell Signaling Tech, cat#:8242) in PBST (1:250) overnight. TBST containing antibody media was removed, washed with 2x500uL TBST/5 minutes each, and stained with DyLight488 anti-rabbit IgG (DI-1488, Vector Laboratories) (1:2000, 1 hours under aluminum foil wrap). After an hour, antibody was removed, washed with 5x500uL TBST, 15 minutes each and finally EZSlide was dismantled. Resulting glass slides was counterstained with Vectashield Antifade Mounting Medium with DAPI (Vector Laboratories, cat#: EW-93952-24), covered with glass coverslips (VWR, cat#48393-081), and stored in a slide-holder packet at 4°C/dark until examined under microscope. Epifluorescent images taken on an Olympus BX41 microscope with Olympus DP74 camera using CellCens software under 20x objective and 10x ocular magnification (200x). Cells manually counted and assessed as follows: Cells with clear nuclear NF-κβ staining greater than control were counted as “positive”, any other cell was counted as “negative” regardless of signal intensity outside of nucleus.

### Western blot analysis

Human (HCH-1) or murine EOC (HGS-1 and HGS-3) cell-lines were seeded in 60mm^3^ (Nest, cat#7050010) overnight in complete DMEM media. Media in 60mm^3^ dishes were replaced with serum free DMEM and stimulated with human IL1β (10ng, 30-45 minutes) or murine IL1β (10ng, 30-45 minutes) without or with pretreatment with UR241-2 (HCH-1, 5-15nM, 4 hrs). For murine EOC cell-lines HGS-1/3, 20, 50 and 150nMoles of UR241-2 were used (Fig-4H). The dishes containing the HCH-1 and HGS-1/3 cells were washed with PBS and lysed with 120uL 1X cell-lysis buffer (Cell Lysis Buffer (10X), Cell Signaling Tech, cat #9803). The lysate was collected in 1.5 ml tube and incubated on ice for 10 min. The lysates were then centrifuged at 14,000 rpm for 10 min at 4°C for removing cell debris. The lysates were electrophoresed using NuPage 4-12% Bis-Tris gels (Invitrogen, cat#NP0322BOX) and transferred semi-dry to PVDF membrane cat#1703966) were all incubated in NuPAGE transfer buffer for 10 min before assembly. The transfer was done at room temperature for 90 minutes at 120 V constant voltage using the PVDF membrane (BioRad, cat#1620177) and filter pads (Bio-Rad, cat#1703969). The PVDF membranes were washed with methanol and then blocked with 5% fat free milk. Chemiluminescent detection was done with a Super Signal West FemtoLuninol/Enhancer (ThermoScientific, cat#1859022 and peroxide cat#1859023) or using Amersham ECL Prime (peroxide solution, cat#RPN2232V2 and enhancer cat#29018903; Cytiva) and probed with p-IRAK4 antibodies (Cell Signaling, cat#11927) (dilution: 1:500). Membranes were imaged using BioRad Chemidoc MP imaging system.

### Colony Formation Assay (Fig-5A)

OVCAR-3, OVCAR-8, and SKOV-3 human EOC cell-lines (20,000/well) were seeded in sterile 8-well glass Millicell EZSlide chambers (Merck Millipore; cat#: PEZGS0816) and cultured in DMEM (Gibco, cat#: 11885-084) supplemented with 10% FBS (Avantor, cat# 89510-186), 1% penicillin/streptomycin (Gibco, cat#:15140-122). Cells were treated with either 5μM, 10μM, or 20μM of UR241-2, or DMSO vehicle for 7 days prior to assessment of colony formation. OVCAR-3 colonies were monitored for 14 days. Phase-contrast images were taken on an Olympus BX41 microscope with Olympus DP74 camera using CellCens software under 20x objective and 10x ocular magnification (200x). Colony borders manually traced in Fiji ImageJ software and the area was measured in μm². Differences between control and treatment groups were analyzed using T-test.

### Cell proliferation assay (Sulforhodamine B Assay, Fig-5C)

OVCAR-3, OVCAR-8, SKOV-3, ES2, and HCH1 human EOC cells (5000/well) were cultured in flat-bottom 96-well plates with 100μL of DMEM (Gibco, cat#: 11885-084) supplemented with 10% FBS (Avantor, cat# 89510-186), 1% penicillin/streptomycin (Gibco, cat#: 15140-122) until ∼50% cell density was met. Cells were then treated for 48 hours with a series of 2-fold dilutions of UR241-2 resulting in the concentrations of 50μM, 25μM, 12.5μM, 6.25μM, 3.12μM, and 1.56μM, as well as DMSO vehicle and negative controls. Cells were fixed with the addition of 70μL 20% TCA and incubated for 1 hour at 4°C. The TCA solution was aspirated, and the plate gently washed once with 200μL dH₂O per well. 100μL of 1x SRB solution diluted in non-supplemented RPMI1640 was added to each well and incubated for 15 minutes at room temperature and protected from light. Solution was aspirated, and plate washed with 200μL 1% acetic acid for 5 minutes on a plate rocker before aspirating off solution. This wash was repeated twice again, and the plate allowed to sit uncovered for residual acetic acid solution to evaporate. 100μL of 10mM Tris base solution was added to each well and the plate was incubated at room temperature for 10 minutes, rocking gently and protected from light. Plate was tapped gently to homogenize dye solution in wells and then read for absorbance at 510nm using the BioTek Synergy 2 plate reader.

### Cell division analyses (Fig-5D)

OVCAR-3 and SKOV-3 cells (10,000 cells/well) were seeded in sterile 8-well glass Millicell EZSlide chambers (Merck Millipore; cat#: PEZGS0816) and cultured in DMEM (Gibco, cat#: 11885-084) supplemented with 10% FBS (Avantor# 89510-186), 1% penicillin/streptomycin (Gibco#: 15140-122). Cells were treated with either 10μM or 20μM of UR241-2, or DMSO vehicle for 48 hours prior to assessment. Cells were then fixed with neutral formalin solution for 25 minutes at 4°C, washed with TBST (3x500mL, 5 minutes each) and stained with p-Histone H2A primary antibody (Cell Signaling Tech, cat#:9718p, 1:1000 in TBST) and incubated overnight. TBST containing antibody was removed, washed 2x500uL TBST 2 times/5 minutes each and stained with DyLight488 anti-rabbit IgG (DI-1488, Vector Laboratories) (1:2000, 1 hours under aluminum foil wrap). After an hour, antibody was removed, washed with 5x500uL TBST, 15 minutes each, and finally EZSlide was dismantled. Resulting glass slides was counterstained with Vectashield Antifade Mounting Medium with DAPI (Vector Laboratories, cat#: EW-93952-24) and covered with glass cover-slides (VWR, cat#48393-081) and stored in a slide-holder packet at 4°C/dark till examined under microscope. Epifluorescent images taken using an Olympus BX41 microscope with Olympus DP74 camera using CellCens software under 20x objective and 10x ocular magnification (200x). Cells manually counted and assessed as follows: cells visibly replicating (metaphase, anaphase) were manually counted and assessed as a percent of the total cells in the field. Cells not totally within the border of the image were not counted.

### Xenograft model (Fig-5E)

SKOV-3 cells (1million cells/mice) were implanted subcutaneously in NSG mice in cold serum free DMEM:Matrigel (1:1). After a week, mice with palpable tumors were randomized and treated with either vehicle (200uL/mice, IP, M-F) or UR241-2 (20mg/kg, IP, M-F). Tumor diameter (L) and width(w) were measured using a digital caliper. On 28^th^ day, mice were euthanized, tumors and peripheral blood from each mouse were collected. The tumor weights were measured using a calibrated balance and hematology of the blood was analyzed using ElementHT-5+ CBC machine.

### Hematology

25-30 µL peripheral blood was collected via retromandibular veins from mice in EDTA-tubes. Complete blood counts (CBC) were analyzed using the HT-5 CBC blood analyzer machine using the system set parameters.

### Effect of UR241-2 on murine EOC tumor burden in syngeneic C57BL/6 mice (Fig-6A-B)

C57BL/6 mice were bred in house from the breeders obtained from Jax laboratories or were purchased from Jax Inc. C57BL/6 mice (n=5 each for control and UR241-2) were implanted with HGS-3 mouse HGS-3 EOC cells (3-4.5 million/mice) intraperitoneally. Two days post tumor cell inoculation, mice were treated with either vehicle or UR241-2 (20mg/kg, 100µL, M-F, IP). DMSO solution of UR241-2 (1µL=200uL, 90mg in 450µL DMSO) was dissolved in formulation made of [H2O (300µL)+PEG6000 (40% in water, 150µL)+EtOH(150µL)] which generates a clear solution after short vortexing. Equivalent volume of the solvent was injected in each animal of the vehicle group. Treatment continued till 44^th^ day. After 45 days, mice were bled via mandibular vein and ∼25µL blood was collected for complete blood counts (CBC) and analyzed in EDTA-tubes using ElementHTS+ system. Mice were euthanized soon after and photographed to record the tumors formed at the injury sites. Spleen, omentum and lavages were collected from each mouse. Omentum and spleen weights were recorded and weighed. Spleen was crushed in serum free RPMI1640 medium (Gibco, cat#: 22400-089) and filtered. Tumors/omentum were also crushed similarly. Crushed cells were broken into single cell suspension and analyzed by flow cytometry following the conditions described below.

### Bulk-seq analysis of the tumors (Fig-7)

Total RNA was isolated using the RNeasy Plus Mini Kit (Qiagen, Valencia, CA) per manufacturers recommendations. The total RNA concentration was determined with the NanopDrop 1000 spectrophotometer (NanoDrop, Wilmington, DE) and RNA quality assessed with the Agilent Bioanalyzer (Agilent, Santa Clara, CA). The Illumina Stranded Total Sample Preparation Kit (Illumina, San Diego, CA) was used for next generation sequencing library construction per manufacturer’s protocols. Briefly, ribosomal-depletion was performed on 100ng total RNA followed by RNA fragmentation. First-strand cDNA synthesis was performed with random hexamer priming followed by second-strand cDNA synthesis using dUTP incorporation for strand marking. End repair and 3‘ adenylation were then performed on the double stranded cDNA. Illumina adaptors were ligated to both ends of the cDNA and amplified with PCR primers specific to the adaptor sequences to generate libraries of approximately 200-500bp in size. The amplified libraries were hybridized to the Illumina flow cell and sequenced using the NovaSeq6000 sequencer (Illumina, San Diego, CA). Paired-end reads of 150nt were generated and a read depth of ∼50M reads was targeted for each sample. Raw reads generated from the Illumina basecalls were demultiplexed using bcl2fastq version 2.19.0. Quality filtering and adapter removal are performed using FastP version 0.20.1 with the following parameters: ““--length_required 35 --cut_front_window_size 1 --cut_front_mean_quality 13 --cut_front --cut_tail_window_size 1 --cut_tail_mean_quality 13 --cut_tail -y -r”. Processed/cleaned reads were then mapped to the mouse reference genome GRCm39 using STAR_2.7.6a with the following parameters: “—twopass Mode Basic --runMode alignReads --outSAMtype BAM SortedByCoordinate – outSAMstrandField intronMotif --outFilterIntronMotifs RemoveNoncanonical –outReads UnmappedFastx”. Genelevel read quantification was derived using the subread-2.0.1 package (featureCounts) with a M31 GTF annotation file and the following parameters: “-s 2 -t exon -g gene_name”. Differential expression analysis was performed using DESeq2-1.28.1 with a P-value threshold of 0.05 within R version 4.0.2 (https://www.R-project.org/). A PCA plot was created within R using the pcaExplorer to measure sample expression variance. Heatmaps were generated using the pheatmap package were given the rLog transformed expression values. Gene ontology analyses were performed using the EnrichR package.

### shRNA knockdown of IRAK4 in HGS-3 murine EOC cells

HGS-3 cells were transfected with murine shRNA plasmid (Santa Cruz Biotechnology, cat# sc-45401-SH) or the control plasmid (control shRNA Plasmid-A, cat#sc-108060) each using lipofectamine (Invitrogen, cat# L3000008) The media was changed after 1 day and cells were treated with increasing doses of puromycin antibiotic (2, 4, 6, 8 and finally 10µM). The surviving cells were collected and a portion of them were frozen in liquid nitrogen. The remaining cells of both control plasmid and IRAK4^KD^ plasmid groups were lysed and expression of IRAK4 was examined by western blot following the methods previously described (IRAK4 antibody, Cell Signaling Technology cat#4363, dilution 1:1000). The same membrane was stripped and probed with GAPDH (Cell Signaling Tech, cat#2118, dilution 1:10,000). The membranes were stripped again and probed with E-cadherin antibody (Cell Signaling Tech, cat #3195, dilution 1:500).

Relative survival of the mouse was measured by implanting the IRAK4kd clone-c2 (7 million/mice, IP, n=5) juxtaposed to Null vector (7 million/mice, IP, n=5). Mice were provided with regular husbandry. The death days were recorded. Survival from implant was estimated using the Kaplan-Meier method. Animals who had not died by the date of analysis at 70 days post implantation were censored. The 2-sided logrank test was used to compare survival between groups.

### Cell cycle analysis

The cell cycle analysis of the HGS-3 null vector clone and IRAK4kd clones were analyzed using flow cytometry instrumentation available with flowcytometry core facilities at the University of Rochester.

### Cell adhesion assay (Fig-7H)

Collagen-coated 96-well plates were prepared by diluting mouse collagen IV (R&D Systems, 3410-010-01 in sterile, cell culture-grade water to a concentration of 10μg/mL. 35μL of collagen was added to each well of a 96-well plate and allowed to dry in the hood under UV light for 2 hours. Each well was washed twice with wash buffer, (0.1% BSA in DPBS), blocked with 100μL 2% BSA in DPBS for 1 hour, followed by two more washes. A plate was prepared for each of the two timepoints: 30 and 60 minutes, respectively. 1 x 10⁶ HGS-3 cells were seeded in a 10cm dish for each concentration tested in complete DMEM (Gibco, cat#: 11885-084) supplemented with 10% FBS (Avantor# 89510-186), 1% penicillin/streptomycin (Gibco#: 15140-122). After allowing the cells to adhere overnight, each respective dish was treated with UR241-2 or DMSO vehicle (Vx) and incubated for 24 hours. Treated cells were harvested and washed twice in serum-free media before being counted and seeded into collagen-coated plates at 2000 cells/well in 100μL serum-free media. At each timepoint, each well was gently washed with wash buffer to remove non-adherent cells, leaving 100μL clean wash buffer in each well. Cells fixed by adding 100μL 10% TCA to each well without discarding existing supernatant and placed at 4°C for one hour before proceeding with SRB protocol.

### Cell migration/wound-healing assay (Fig-7I)

HGS-3 cell line was seeded in a 24-well plate, with 10,000 cells per well in complete DMEM (Gibco, cat#: 11885-084) supplemented with 10% FBS (Avantor# 89510-186), 1% penicillin/streptomycin (Gibco#: 15140-122). Once cells approached ∼70% cell density, they were treated with UR241-2 for 24 hours. After treatment, each well received a single vertical scratch using a P1000 tip. Scratched wells were rinsed once with media to remove detached cells and given fresh media. Initial images were taken immediately following this, represented by the “0” timepoint. Images were taken of scratch at the same location in 12-hour intervals for 24 hours. Images were taken on an Olympus CKX41 inverted phase-contrast microscope with an Olympus DP74 camera and CellSens software. Image analysis was completed using Fiji’s ImageJ software and the wound healing size tool plugin developed by Suarez-Arnedo et al^60^.

### Statistical analysis

Statistical significance between independent groups for non-repeated measurements was evaluated using the Wilcoxon-Mann-Whitney test, a nonparametric T-test. This method applies to outcomes measured on a continuous scale when small samples may hamper confirmation of distributional assumptions. Xenograft model data was analyzed using a repeated measures analysis of variance. The statistical model was fit using maximum likelihood estimation with treatment, day, and the interaction between treatment and day as fixed effects. Mouse was considered a random effect, and the correlation of repeated measures on the same mouse over time was handled using a first-order ante-dependence variance-covariance structure which varied by treatment. Model assumptions were verified graphically. If the Type 3 F-test for the interaction between treatment and day was significant, then treatment comparisons at each day were conducted using a T-test. Analysis was conducted using SAS v9.4 Proc Mixed (Cary, NC).

## Data availability

Raw data for Figures 1-6 has been provided in an Excel spreadsheet. The raw sequencing data of the vehicle or UR241-2 treated tumors (Fig-7) were deposited to the GEO (accession code: GSE296588). Data will be shared on reasonable requests. Data access requests should be submitted to RKS. Requests will be reviewed by University of Rochester and Empire Discovery Institute jointly and responses will be provided within approximately 8-10 weeks. Approved users must comply with data use agreements specifying permitted use and restrictions.

## Acknowledgement

Empire Discovery Institute (EDI) paid for Pan-kinase screening and ADMET studies. EDI played no role in writing this manuscript. Mechanistic model (Model-1) depicting the mechanism of action of UR241-2 was drawn using Biorender software licensed to the University of Rochester. The authors would like to thank the University of Rochester Flow Cytometry Core and Genomics Core for their assistance. ChatGPT was used only for grammar refinement of author-written content, with no role in study design, drafting of the manuscript, data analysis, or interpretation.

## Author’s contribution

JPM performed flow cytometry and edited the manuscript. NK performed western blotting. CWS performed cell adhesion and migration assays, microscopy, and analyzed/quantified images. NAS performed docking and simulations. EL performed animal experiments and did the treatments, measured tumor sizes and animal weights and performed CBC analysis repeatedly. MS performed the statistical analysis. KKK helped develop IRAK4^KD^ HGS-3 cell clones and performed NF-kβ reporter assay. MA and RK designed and partially funded ADME assays. SS analyzed the flow data under supervision of JB. CMA provided OVCAR-3-NF-κ reporter cell-line and methods of use. JA, CB and EP performed bulk-seq. MEB drew the model-1. MW edited the manuscript. MKK conducted the synthesis of UR241-2 and analogs. RRT, RGM, LMC provided laboratory resources, funding and critically read and edited the manuscript multiple rounds. MWB provided KO animals, guidance and reviewed the data. RKS brought resources with RGM and MWB, conceived the study, designed and synthesized UR241-2 first round and wrote the manuscript.

## Ethics statement

Animal experiments were performed with prior approval of University Committee of the Animal Resources (UCAR) of the University of Rochester. Approval number (UCAR:2016-011E)

---

**Model-1:**
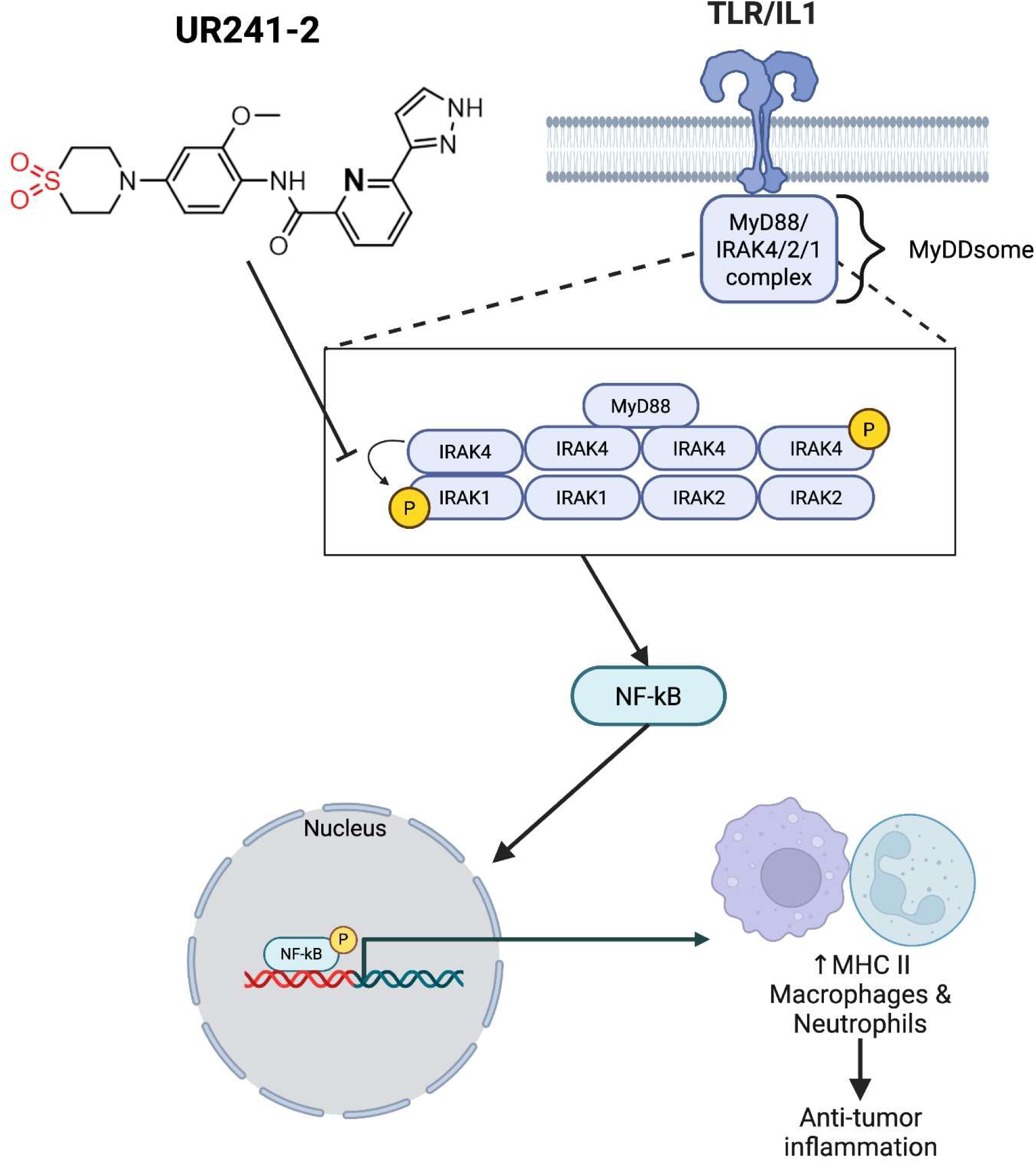
Based on the data, it is proposed that IRAK4 kinase inhibition by UR241-2 led to blocked activity of NF-κβ resulting in the activation of MHCII macrophages and neutrophils which suppressed tumor growth at the site of injury. Model was prepared using Biorender software.

## Supplementary Information

**Supplementary Figure-1:**
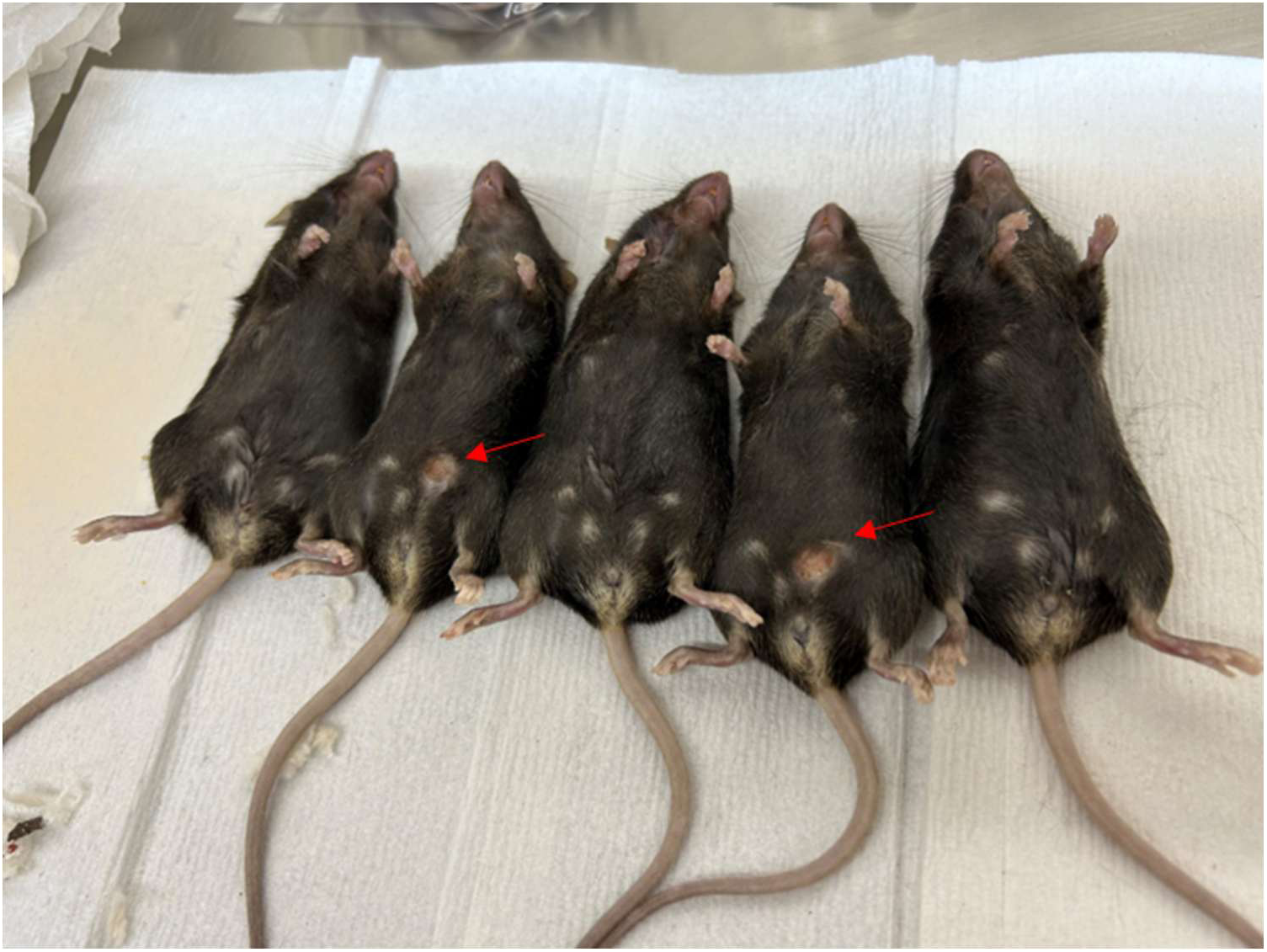
HGS-3 murine high-grade serous EOC cells (3-4.5 million/per mice) were implanted intraperitoneally using 21-gauge needle in C57BL/6 WT and C57BL/6 Nlrp3^KO^ mice. Mice were observed for 45-50 days and euthanized. Tumors formed on needle injury site, protruding at the skin as well as in the peritoneum and on the omentum, shown by red arrows, were isolated, weighed and frozen in liquid -nitrogen. Lavages via washing with sterile PBS(5mL) were also collected. The studies were repeated twice. A representative experiment is shown. Weights of the omental did not differ between C57BL/6 WT (Figure-1H) and Nlrp3^KO^ mice. Similarly, the tumor sizes at the site of injury did not differ between C57BL/6 WT and Nlrp3^KO^ mice.

**Supplementary Figure-2:**
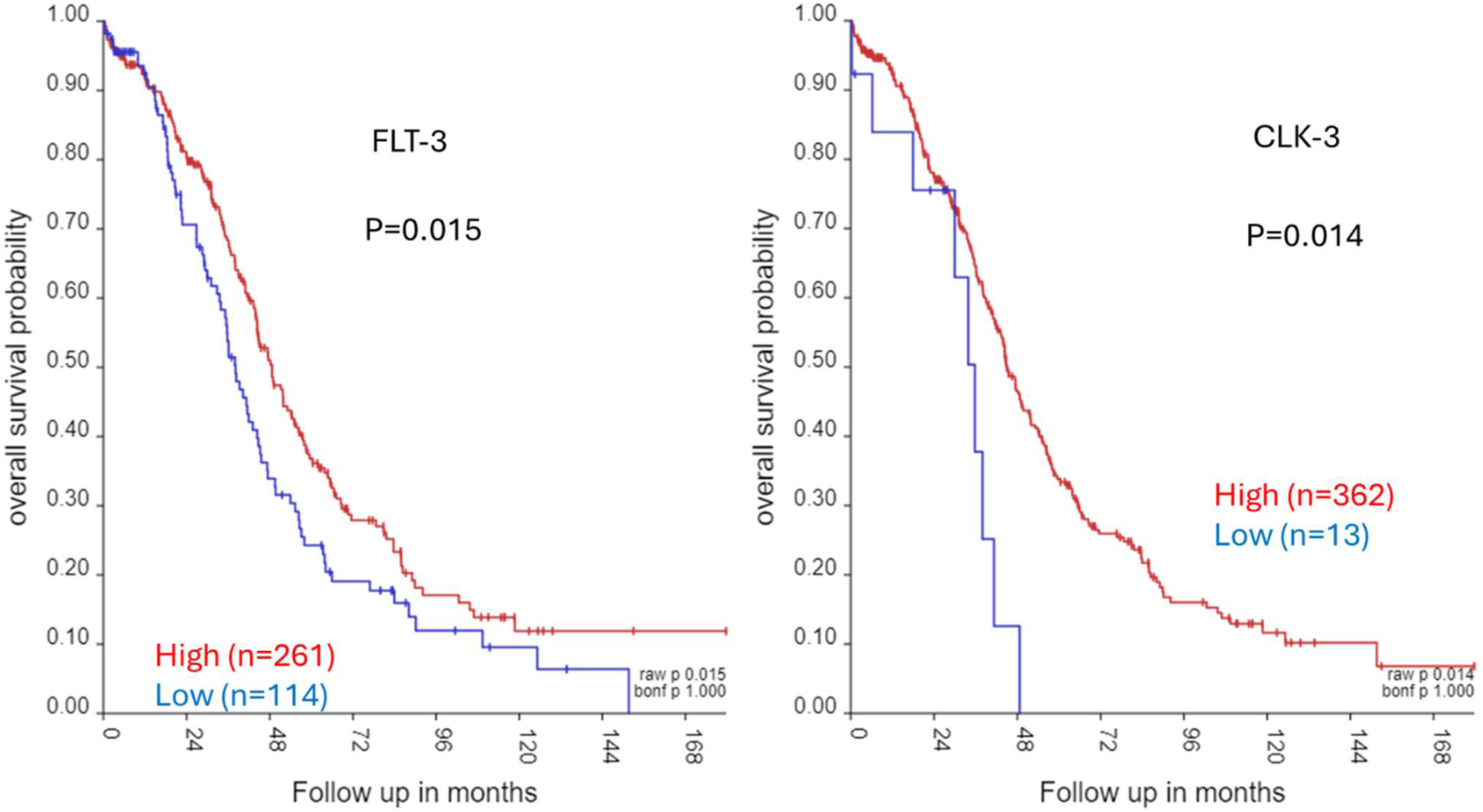
Analysis of ovarian serous cystadenocarcinoma (2022-v32) microarray data (TCGA-381-tpm-gencode36) of ovarian cancer patients using R2 Genomics Analysis and Visualization Platform tools showed that FLT-3 and CLK-3 mRNA overexpression predicts poor survival.

**Supplementary Figure-3:**
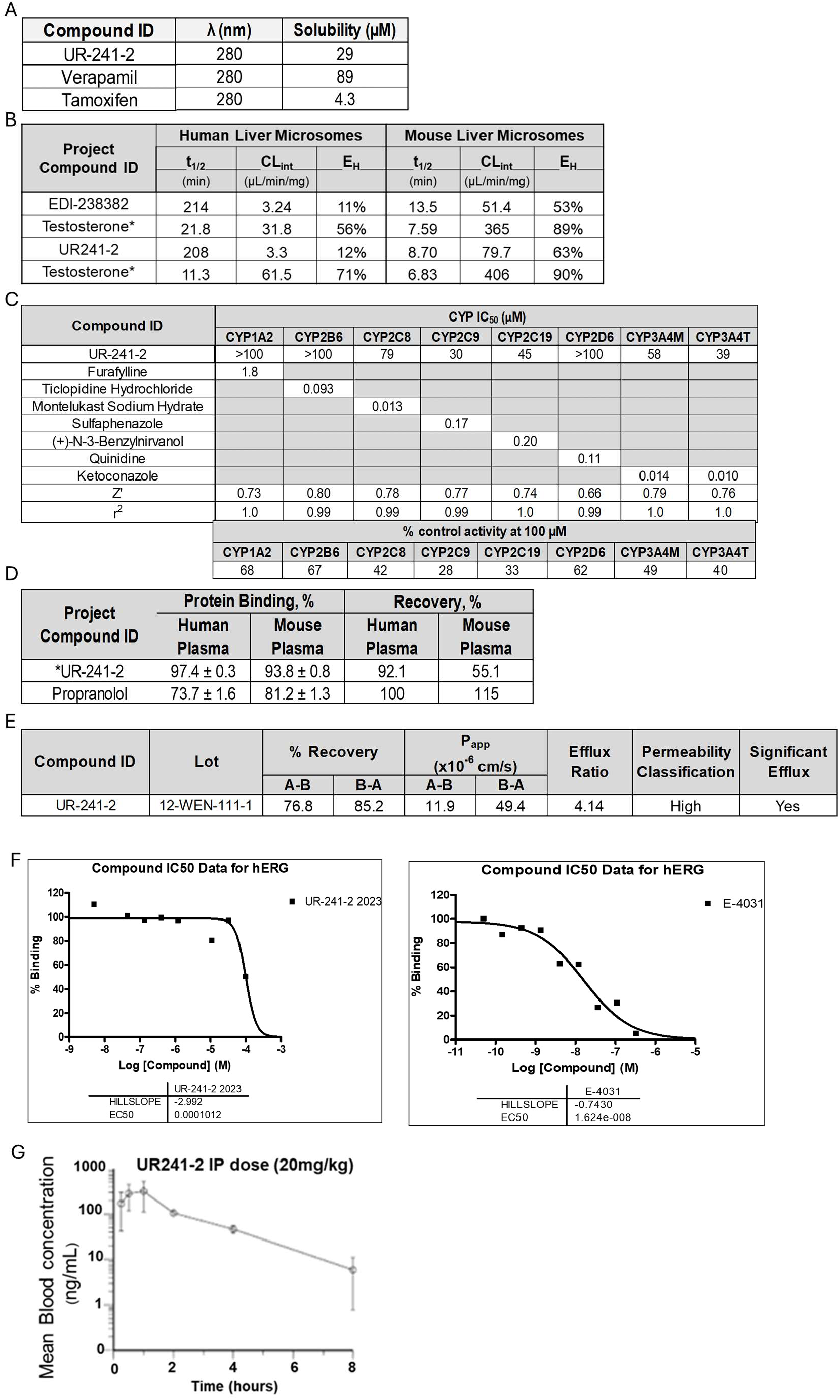
(**A**): Kinetic solubility of UR241-1 is shown. (**B**): Quantified stability of UR241-2 in human and mouse liver microsomes is shown. (**C**): Inhibition of CYP450 isoforms by UR241-2 is quantified using HPLC. Controls used were furafyllin, Ticlopidine HCL, Montelukast sodium hydrate, sulfaphenazole, N-3-benznirvanol, Qunidine, ketoconazole. CYP isoforms affected by UR241-2 in terms of %-inhibition are shown. (**D**): Human and murine plasma protein binding of UR241-1 is shown in % units. Propranolol was used as a control. (**E**): CaCo-2 cell permeability of UR241-2 is shown. Efflux ratio of 4.14 indicates that UR241-2 faces significant efflux. (**F**): UR241-2 does not inhibit hERG. IC_50_ is 101.2µM. E-4031 was used as control. IC_50_ for E-4031 was 1.62e-^08^. (**G**): Pharmacokinetic (PK) of UR241-2 at 20mg/kg administered IP is shown. It is shown that ∼8ng/ml concentration of UR241-2 is maintained up until 8^th^ hour of monitoring.

**Supplementary Figure-4:**
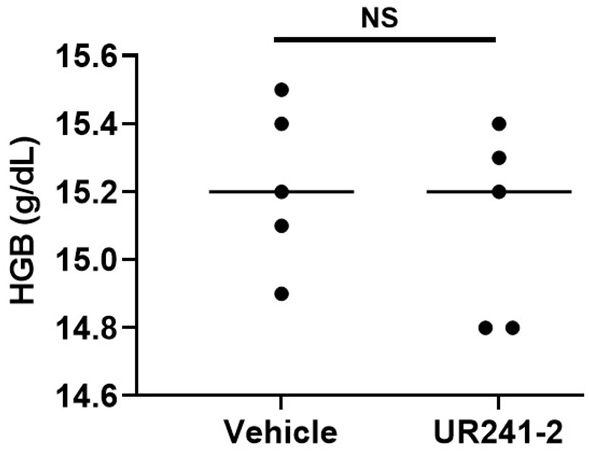
Analysis of the peripheral blood showed that hemoglobin (HB) levels did not differ between the vehicle and UR241-2 treated mice.

**Supplementary data-5:**
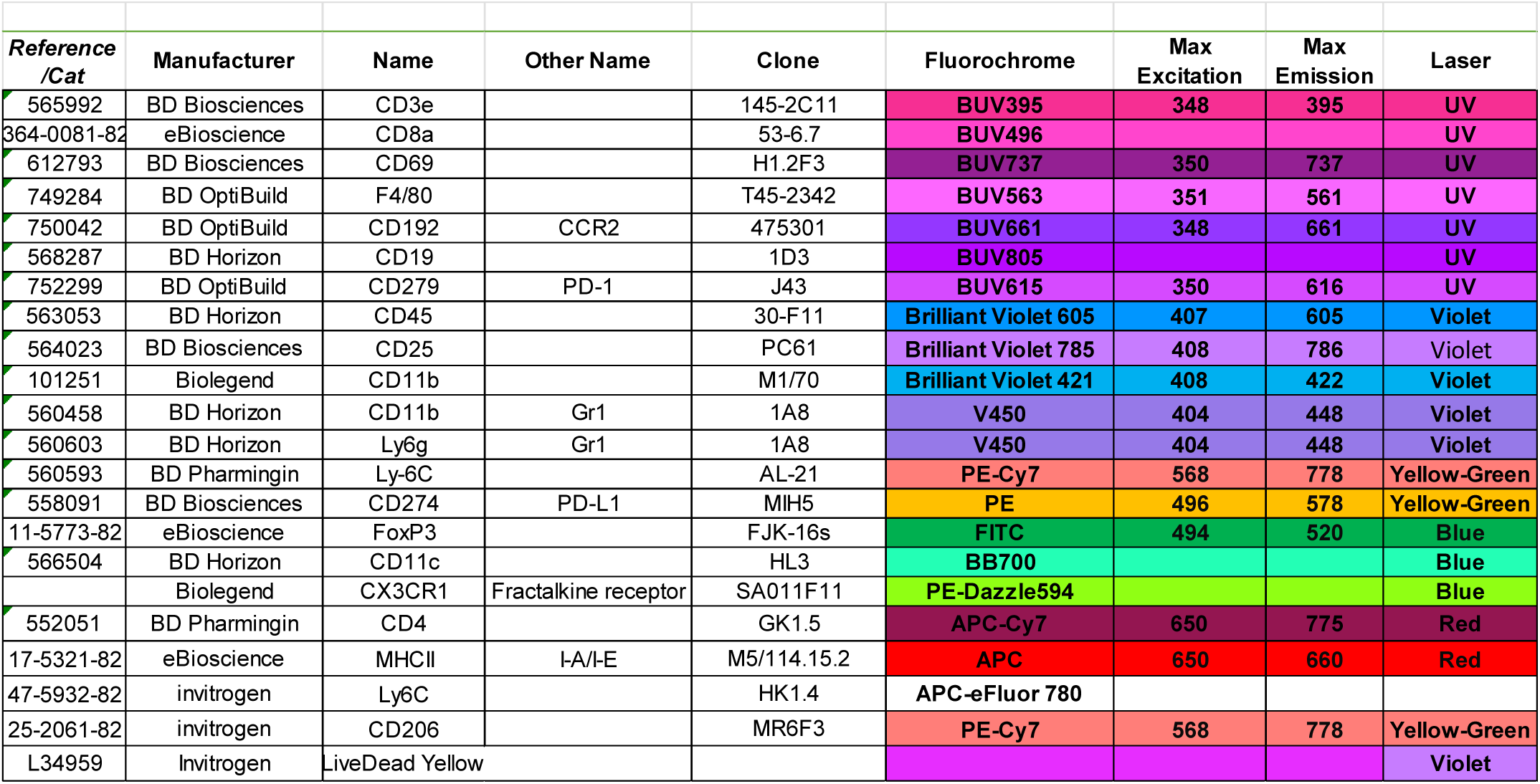
List and catalog details (reference no, manufacturer, name, other name, Clone, fluorochrome, max and min excitation, laser) of flow cytometry antibodies used in this study.

**Supplementary Data-6**

**NMR and Mass spectrometry data for UR241-2:**

^1^H NMR (400 MHz, CDCl_3_), δ: 10.52 (s, 1H), 8.51 (d, *J* = 8.7 Hz, 1H), 8.21 (d, *J* = 6.6 Hz, 1H), 8.04 – 7.93 (m, 2H), 7.72 (d, *J* = 2.1 Hz, 1H), 6.98 (d, *J* = 2.1 Hz, 1H), 6.60 (dd, *J* = 8.8, 2.5 Hz, 1H), 6.56 (d, *J* = 2.4 Hz, 1H), 4.01 (s, 3H), 3.86 – 3.77 (m, 4H), 3.21 – 3.11 (m, 4H). LCMS (m/z): 428.3 [M+H] ^+^.

**NMR and Mass spectrometry data for PSP-0099:**

^1^H NMR (400 MHz, CDCl_3_), δ: 10.51 (s, 1H), 8.48 (d, J = 9.3 Hz, 1H), 8.21 (d, J = 7.2 Hz, 1H), 8.02 – 7.92 (m, 2H), 7.72 (d, J = 1.9 Hz, 1H), 6.98 (s, 1H), 6.60 (d, J = 6.5 Hz, 2H), 4.00 (s, 3H), 3.57 – 3.50 (m, 3H), 2.86 – 2.77 (m, 4H). LCMS (m/z): 396.3 [M+H] ^+^.

**NMR and Mass spectrometry data for PSP-0100:**

^1^H NMR (400 MHz, CDCl_3_), δ: 10.51 (s, 1H), 8.49 (d, J = 8.7 Hz, 1H), 8.21 (d, J = 7.3 Hz, 1H), 7.97 (dt, J = 15.3, 7.5 Hz, 2H), 7.72 (d, J = 1.5 Hz, 1H), 6.98 (s, 1H), 6.64 (dd, J = 8.8, 2.4 Hz, 1H), 6.60 (d, J = 2.4 Hz, 1H), 3.99 (d, J = 10.2 Hz, 3H), 3.95 (dd, J = 10.2, 3.2 Hz, 2H), 3.54 (dd, J = 18.1, 3.7 Hz, 2H), 3.00 – 2.85 (m, 4H). LCMS (m/z): 412.2 [M+H] ^+^.

**Supplementary data-7.**
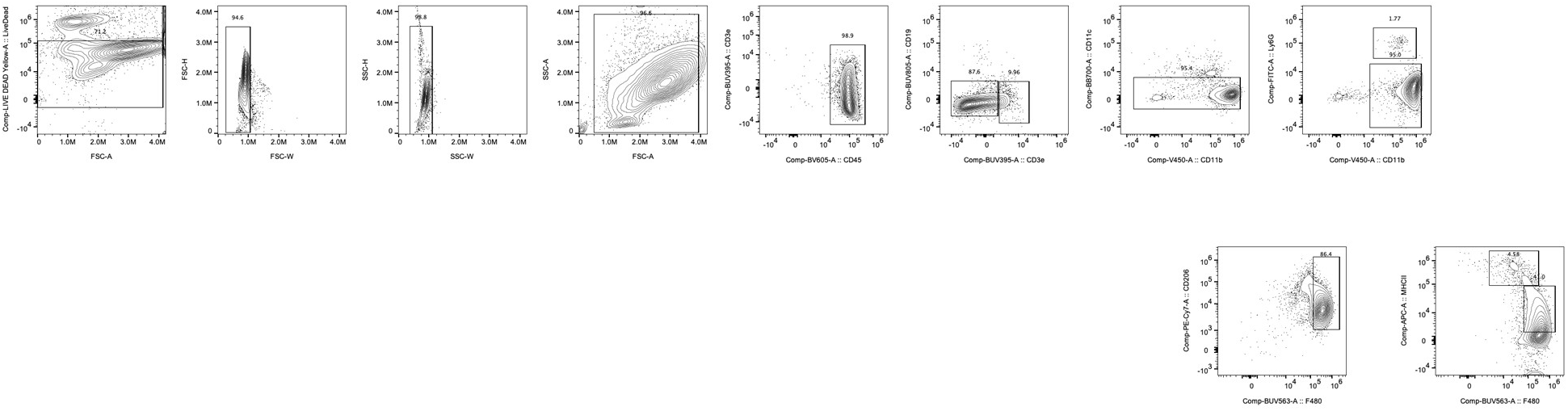
Gating strategy.

